# Phagocytic predation by the fungivorous amoeba *Protostelium aurantium* targets metal ion and redox homeostasis

**DOI:** 10.1101/690503

**Authors:** Silvia Radosa, Jakob L. Sprague, Renáta Tóth, Thomas Wolf, Marcel Sprenger, Sascha Brunke, Gianni Panagiotou, Jörg Linde, Attila Gácser, Falk Hillmann

**Affiliations:** Junior Research Group *Evolution of Microbial Interaction*, Leibniz Institute for Natural Product Research and Infection Biology - Hans Knöll Institute (HKI), Jena, Germany; Institute of Microbiology, Friedrich Schiller University Jena, Jena, Germany; Department of Microbiology, University of Szeged, Szeged, Hungary; Research Group Systems Biology and Bioinformatics, Leibniz Institute for Natural Product Research and Infection Biology - Hans Knöll Institute (HKI), Jena, Germany; Department of Microbial Pathogenicity Mechanisms, Leibniz Institute for Natural Product Research and Infection Biology - Hans Knöll Institute (HKI), Jena, Germany; Institute of Bacterial Infections and Zoonoses, Federal Research Institute for Animal Health – Friedrich-Löffler-Institute, Jena, Germany

## Abstract

Predatory interactions among microbes are considered to be a major evolutionary driving force for biodiversity and the defense against phagocytic killing. The fungivorous amoeba *Protostelium aurantium* has a wide fungal food spectrum but strongly discriminates among major pathogenic members of the *Saccharomycotina*. While *C. albicans* is not recognized, *C. glabrata* is rapidly internalized, but remains undigested. Phagocytic killing and feeding by *P. aurantium* is highly effective for the third major fungal pathogen, *C. parapsilosis.* Here we show that the different prey patterns of the three yeasts were reflected by distinct transcriptional responses, indicating fungal copper and redox homeostasis as primary targets during intracellular killing of *C. parapsilosis*. Gene deletions in this fungus for the highly expressed copper exporter Crp1 and the peroxiredoxin Prx1 confirmed their role in copper and redox homeostasis, respectively and identified methionine biosynthesis as a ROS sensitive metabolic target during predation. Both, intact Cu export and redox homeostasis contributed to the survival of *C. parapsilosis* not only when encountering *P. aurantium*, but also in the presence of human macrophages. As both genes were found to be widely conserved within the entire *Candida* clade, our results suggest that they could be part of a basic tool-kit to survive phagocytic attacks by environmental predators.

## Introduction

Members of the genus *Candida* are among the leading causative agents of fungal infections worldwide with *Candida albicans* being responsible for the majority of candidiasis cases, followed by *C. glabrata* and *C. parapsilosis* (1). All three *Candida* species are known to be commensals and are frequently residing in oral cavities, the gastrointestinal tract, vaginal mucosa or on the skin. Environmental reservoirs for any of these species have rarely been documented, but recent isolations of *C. parapsilosis* or *C. albicans* from pine and oak trees, respectively, suggest that these might exist (2–4). *C. glabrata*, in turn, has been enriched from fermented foods and grape juice (5, 6). Within the human host, all three are able to counteract the phagocytic attacks of macrophages and neutrophilic granulocytes to some extent, using different strategies and molecular tool-kits (7).

An outer layer of mannoproteins masks pathogen-associated molecular patterns (PAMPs) on the surface of *C. albicans*, thus hindering the initial recognition of the fungus *via* cell wall β-glucans (8). Even after its ingestion, *C. albicans* can escape from innate immune phagocytes by hyphae formation which triggers the cytolytic death of the host cell (9–11). *C. parapsilosis* is also able to survive in the restricted phagosomal environment and forms pseudohyphae after its internalization by macrophages (12). However, its rates of ingestion and killing by neutrophils and macrophages were reported to be higher than for *C. albicans* (12–14). Intracellular filamentation, in turn, is not observed for *C. glabrata*, which instead can survive and even replicate in the yeast form inside modified phagosomal compartments of macrophages (15, 16).

Estimates indicate that *C. glabrata* may be separated from the other two *Candida* species by more than 300 million years (17), well before their establishment as commensals. Comparative genome analysis of *C. glabrata* and its closest relatives have suggested that adaptations preceding its commensal stage may have facilitated traits that later enabled pathogenicity (18–20). Fungal infections originating from species even without any clear history of commensalism have further raised questions on the role of environmental factors as early promoters of virulence-associated traits.

Predator-prey interactions are considered as drivers of an evolutionary arms race and occur frequently, even among microbes. Humans and higher animals are indirectly affected, as some microbial defenses against phagocytic predators are thought to be also effective against innate immune cells such as macrophages and neutrophilic granulocytes. These trained defenses may have favored certain microbes to establish commensalism or appear as new pathogens (21). Experimental studies have corroborated this idea using well-known model amoebae such as *Dictyostelium discoideum* or *Acanthamoeba castellanii* (22–26). *Protostelium aurantium* is another representative of a wide-spread group of amoebae with a fungivorous life-style (27–31). The amoeba was recently found to feed on a wide range of basidiomycete and ascomycete yeast species, with *C. parapsilosis* being the most efficient food source, while *C. albicans* and *C. glabrata* escaped the predation at the stage of recognition or intracellular processing, respectively (32). In this study, we investigated the responses of *C. albicans, C. glabrata*, and *C. parapsilosis* when confronted with the fungivorous predator. Our findings demonstrate that copper and redox homeostasis are central targets during phagocytic predation by *P. aurantium* and suggest that such basic anti-phagocytic defense strategies may have been trained during an arms race with an environmental predator.

## Results

### The predation responses of the three *Candida* species reflect their different prey patterns

While *C. albicans* and *C. glabrata* can escape *P. aurantium* at the stage of recognition or intracellular processing, respectively, *C. parapsilosis* serves as a highly efficient food source (32). To elucidate common, as well as species-specific reactions to the presence of the predator, we conducted high-throughput RNA sequencing of each of the three *Candida* species in co-cultures with *P. aurantium.* Yeast cells were confronted with trophozoites of *P. aurantium* for 30 and 60 minutes prior to sampling for RNA isolation. For *C. parapsilosis*, a total of 667 genes were upregulated (log_2_FC > 1.5 with p<0.01 according to EdgeR), and 588 genes were downregulated (log_2_FC <-1.5, p<0.01), while in *C. albicans*, a total of 903 genes were upregulated, and 937 genes were downregulated at both time points (Fig. 1A). In *C. glabrata*, 1218 genes were upregulated, while only 273 genes were found to be downregulated (Fig. 1A). A complete list of DEGs for each species and time point is given in Dataset S1.

**Fig. 1:**
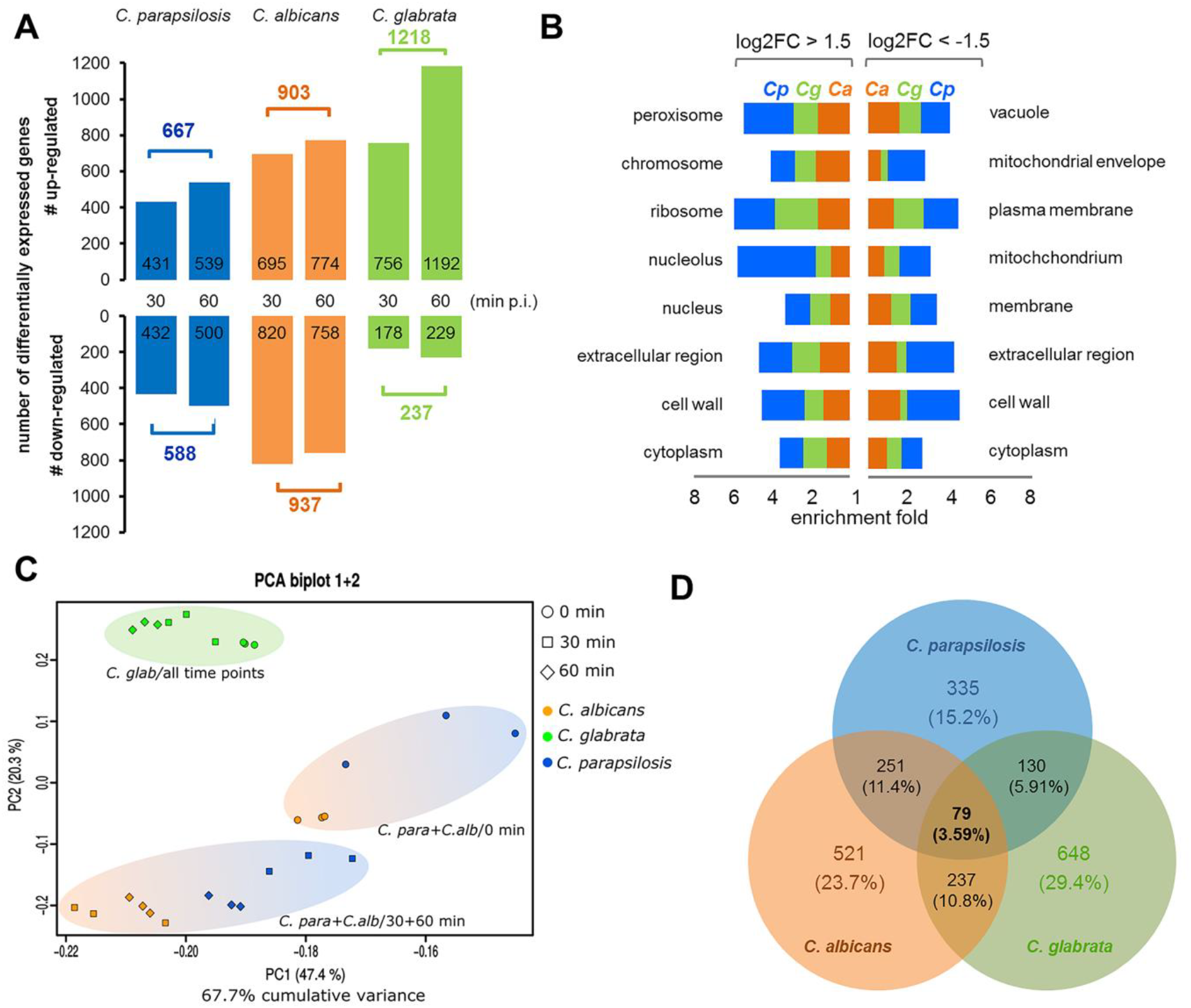
Differential gene expression in *C. parapsilosis, C. albicans*, and *C. glabrata* in response to *P. aurantium.* **(A)** Total numbers of differentially expressed genes (DEGs) of *Candida* spp. in the presence of *P. aurantium* after 30 and 60 min. Genes were considered as differentially expressed when the log_2_ fold-change in the transcript level was ≥1.5 or ≤-1.5 and p≤0.01 according to EdgeR at either of the two time points. **(B)** Gene ontology (GO) clusters for cellular components and their enrichment in up- (left panel) and downregulated (right panel) genes of *C. parapsilosis* (Cp, blue), *C. glabrata* (Cg, green), and *C. albicans* (Ca, orange) **(C)** Principle component analysis (PCA) for read count values from all orthologous genes of the three *Candida* species. PC1 and PC2 explain about 68% of the overall variance within the data set and clearly separate all *C. glabrata* samples (green, top left) from those of *C. albicans* (orange) and *C. parapsilosis* (blue). For the two latter species, there is an additional separation between control time-point at 0 min (round) and time points 30/60 min (square/diamond). **(D)** Venn diagram displaying an overlap in the differential expression of all common orthologues at 30 and 60 min. Of all 3,735 orthologues in total, 2,201 were differentially expressed orthologues (DEOs) and 79 were common (3.6%) to all three species.

To address the biological significance of up- and downregulated genes, we analyzed their gene ontology annotations for the enrichment of defined categories in *molecular function, cell component*, and *biological process* (Fig. 1B, Fig. S1, Dataset S2). Overall, the enriched categories for all three *Candida* species partially overlapped, most likely resulting from general metabolic adaptations: e. g. when grouped by *molecular function*, transferase and ligase activity were categories common to all three fungi among the upregulated genes. However, for *C. glabrata*, there was no significantly enriched *biological process* among the downregulated genes (Fig. S1). Also, transporter and kinase activity were the only two *molecular functions*, which were enriched among the downregulated genes of *C. glabrata*. In sharp contrast, transporters were found to be generally upregulated in *C. albicans* and *C. parapsilosis*. Higher expression levels of RNA binding, helicases, and nucleotidyl transferases were unique to *C. parapsilosis*, the preferred prey, implicating that transcription and translation could be most severely affected in this fungus. The finding that the nucleolus and *biological process* categories for RNA metabolism and ribosome biogenesis were all only enriched in *C. parapsilosis* also supports this conclusion. Further, the extracellular region and the cell wall were more severely affected in *C. parapsilosis* than in *C. albicans* or *C. glabrata* (Fig. 1B).

### The core response of *C. albicans, C. parapsilosis*, and *C. glabrata* to the presence of *P. aurantium*

A principal component analysis was used to determine the dynamic variations in the orthologous DEGs. For *C. glabrata*, the expression of orthologous genes in response to amoeba predation was clearly distinguishable from *C. parapsilosis* and *C. albicans*, and also showed less variation between the different time points (Fig. 1C and the cluster dendrogram in Fig. S2). Although more than 1,200 genes were activated in *C. glabrata* overall, only 4 % of the differentially expressed orthologs were unique to the first time-point at 30 min. These numbers clearly differed for *C. parapsilosis* (15 %) and *C. ablicans* (18 %), and thus, displayed more variance over time. For these two, it was also evident that their transcriptional profiles at 30 and 60 min clustered closer together than with any time-point from *C. glabrata*. To identify a common responsive gene set of all three fungi, we compared the differential expression among all their orthologues (DEOs). Of the overall 3,735 orthologous genes among the three species, differential expression was found for 2,201 genes at either one of the two later time points (Fig. 1D). Reflecting the diverse interaction patterns of the three species, only 79 orthologous genes were differentially regulated in all three *Candida* species, representing the core response to the presence of *P. aurantium* (Fig. 1D). Among those, 48 genes were commonly induced and eight genes were commonly repressed in all three species at either of the time points, while 23 genes showed opposite regulation between the species (Fig. 2).

**Fig. 2:**
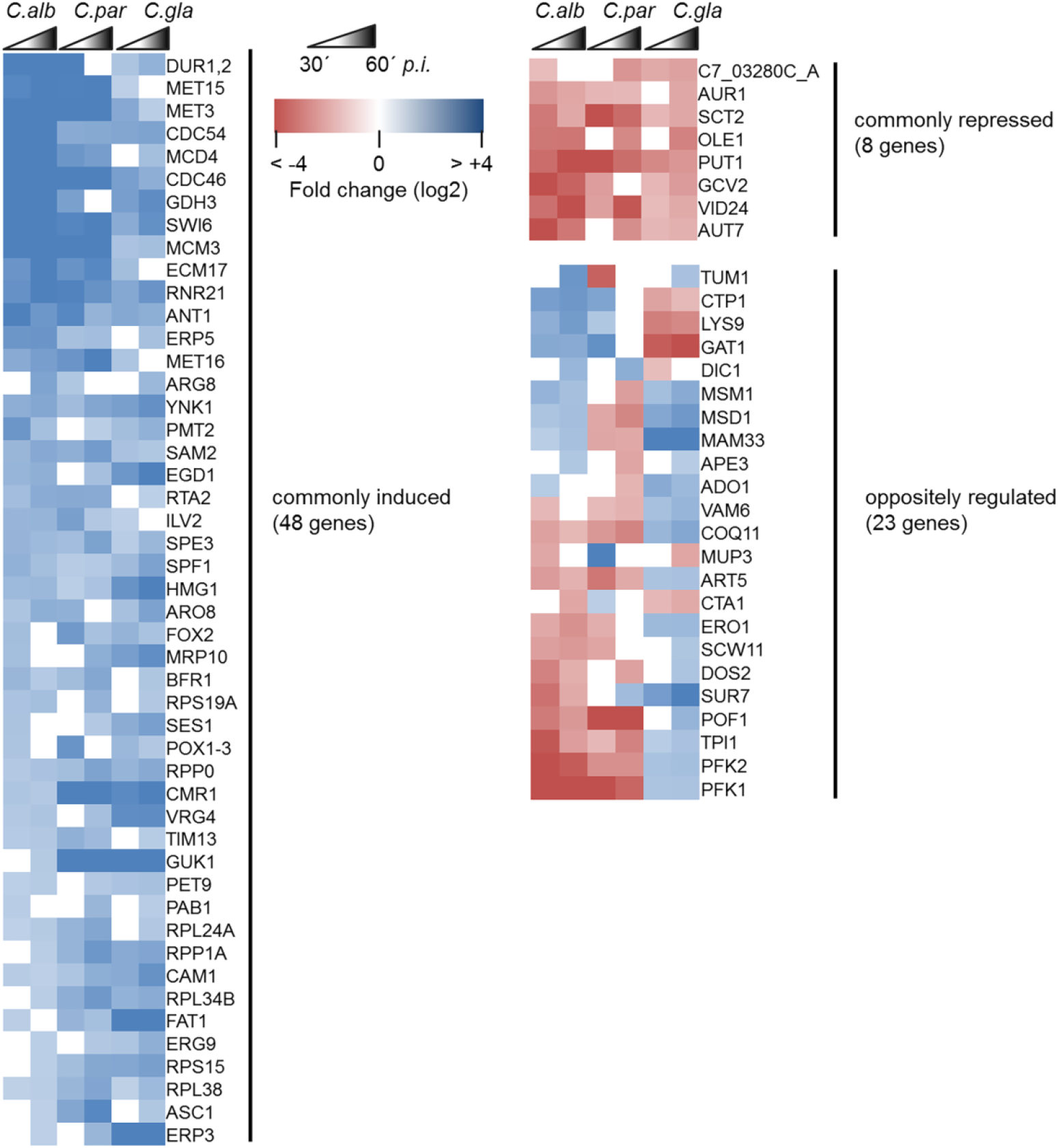
Heat map of expression for all 79 differentially expressed orthologues (DEOs) during the confrontation with *P. aurantium*. All DEOs were grouped according to their transcription profile at 30 min and 60 min *p.i.* in comparison to 0 min, and considered as commonly induced or commonly repressed if they shared the same expression pattern among all three species. DEOs were considered as oppositely regulated if their expression tendency differed in one of the three species. Red and blue colors represent down- and upregulated genes, respectively. Gene names are based on the orthologues of *C. albicans*.

The corresponding sets of genes were further analyzed for shared GO terms in biological processes (Dataset S3). Commonly enriched categories included the sulfur amino acid metabolic process (GO:0000096) comprising genes such as *SAM2, MET3, ECM17* (*MET5*)*, MET15*, and *MET16*; all playing a role in the metabolism of methionine. A plethora of genes, predicted to be involved in organo-nitrogen compound biosynthetic process (GO:1901566) such as the amino acid biosynthesis enzymes *ILV2* and *ARG8*, a P-type calcium-transporting ATPase encoded by *SPF1*, or genes with role in fatty acid beta-oxidation (GO:0006635) like *ANT1, FOX2* or *POX1-3*, were commonly induced as well. The most highly enriched GO term was found to be “negative regulation of helicase activity” comprising three *MCM* genes: *CDC54* (*MCM4*), *CDC46* (*MCM5*), and *MCM3*; all known to be a part of Mcm-complex, necessary for unwinding the DNA double helix and triggering fork progression during DNA replication (33).

Noteworthy is the induction of the *DUR1,2* gene in all three species, encoding the urea amidolyase and previously shown to be important for the survival of *C. albicans* in macrophages (34). No GO category was found to be enriched within the eight commonly downregulated genes. Nevertheless, three out of eight genes, namely *OLE1, SCT2* and *AUR1*, function in lipid biosynthetic processes and most probably play an important role in the integrity of the cell membrane. Interestingly, GO enrichment analysis revealed the “glycolytic process through fructose-6-phosphate” (GO:0061615) as a highly overrepresented category within the oppositely regulated set of genes; all genes annotated to this category, namely *TPI1, PFK1* and *PFK2*, were downregulated in *C. albicans* and *C. parapsilosis*, while in *C. glabrata* they showed an increase in transcript level.

### *P. aurantium* predation targets copper and redox homeostasis in *C. parapsilosis*

Of all three species, *C. parapsilosis* represented the preferential food source for *P. aurantium* and thus, we conducted a deeper characterization of the 1255 DEGs (667 genes with log2FC ≥ 1.5, and 588 genes with log2FC ≤ −1.5) from *C. parapsilosis* using the GO Slim tool which maps DEGs to more general terms and broad categories (35). Most genes were uncharacterized, could not be categorized and were involved in unknown biological processes or mapped to “regulation” as a general category. These were not further analyzed. “Transport” and “stress response” were the two most frequent biological processes and were further selected to search for more specific categories (Fig. 3). The “extra-nuclear transport of ribonucleoproteins” was highly enriched, as could be expected from the results obtained from the general enrichment analysis for *C. parapsilosis*. We further found the transport of transition metal ions to be overrepresented with several genes encoding orthologous proteins for the transport of Fe and Zn being differentially regulated in response to *P. aurantium* (Tables S1 and S2).

**Fig. 3:**
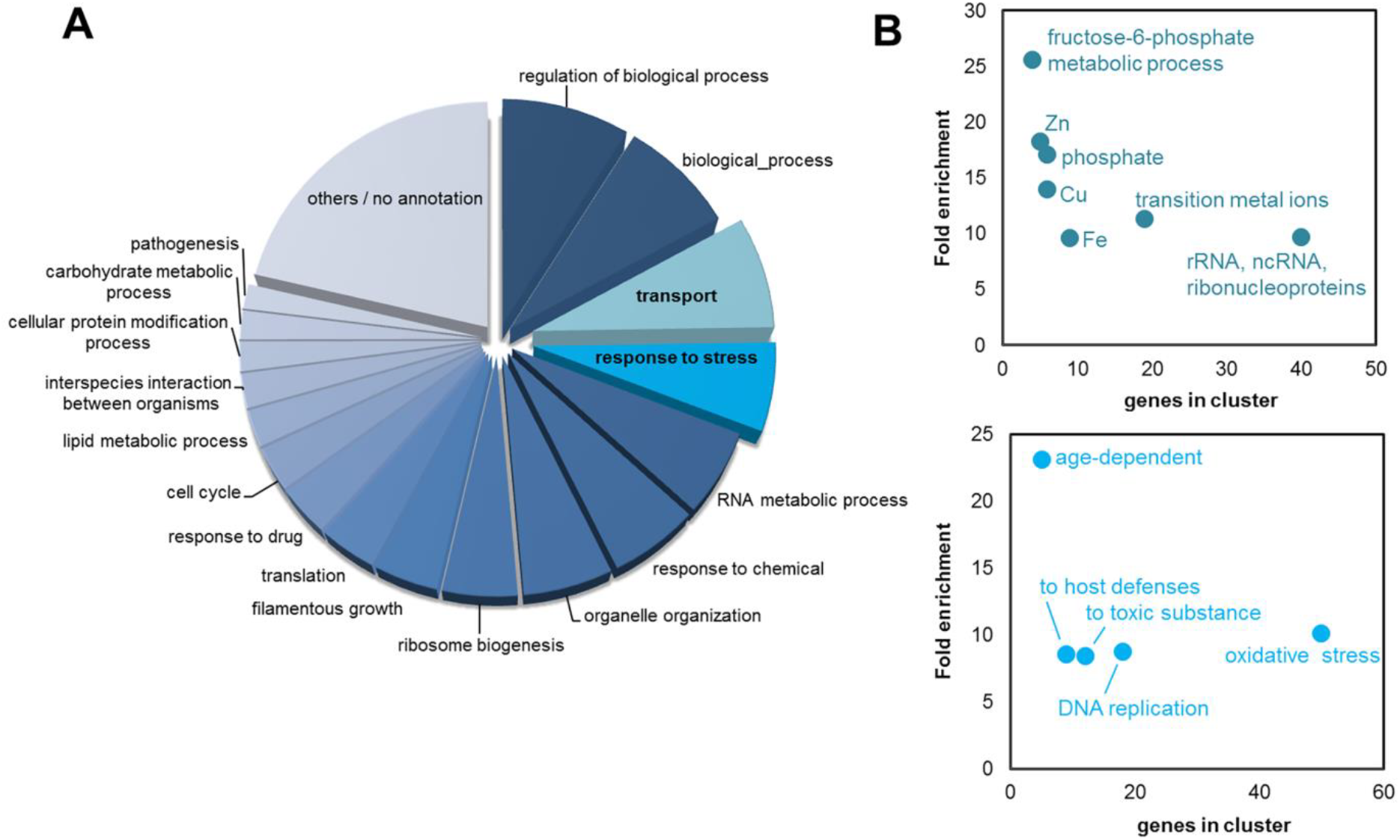
GO Slim categorization of the DEGs from *C. parapsilosis* during the confrontation with *P. aurantium*. **(A)** All 1255 DEGs from 30 min and 60 min after the confrontation with *P. aurantium* were categorized according to GO SLIM processes. **(B)** Genes mapped to the GO SLIM categories “transport” and “response to stress” were further analyzed for more specific GO Terms. Categories with the highest enrichment (top 7 for “transport” and top 5 for “response to stress”) are displayed as a 2D dot plot for the number of genes in the respective cluster and fold enrichment (p-value <0.005).

Two Cu transporters were among the most strongly differentially regulated genes in *C. parapsilosis* (Table 1). The most upregulated gene upon amoeba predation (log_2_FC of approx. 9 at 30 min and 8 at 60 min) was found to be CPAR2_203720. This gene is an orthologue to of *C. albicans CRP1* (*orf19.4784*), encoding a copper-transporting P1-type ATPase, which mediates Cu resistance and is induced by high Cu concentrations (36).

**Table 1:** Expression of *C. parapsilosis* genes involved in the transport of Cu ions (GO:0006825)

| name | log2FC<br>30 min | log2FC<br>60 min | description | *C. albicans* orthologue |
| --- | --- | --- | --- | --- |
| CPAR2_203720 | 9.36 | 8.18 | copper-exporting ATPase activity, role in cadmium ion transport, cellular copper ion homeostasis, copper ion transport, silver ion transport and plasma membrane localization | C1_09250W_A/CRP1 |
| CPAR2_210510 | -2.36 | -2.46 | copper uptake transmembrane transporter activity, role in cellular copper ion homeostasis, copper ion import, intracellular copper ion transport and fungal-type vacuole membrane localization | C1_08620W_A/CTR2 |
| CPAR2_300620 | -7.66 | -8.01 | ferric-chelate reductase activity, role in copper ion import, iron ion transport and plasma membrane localization | C7_00430W_A |
| CPAR2_406100 | -1.81 | -3.67 | copper ion transmembrane transporter activity, inorganic phosphate transmembrane transporter activity and role in cellular copper ion homeostasis, copper ion transmembrane transport, phosphate ion transmembrane transport | C2_09590C_A |
| CPAR2_701290 | -3.09 | -3.26 | copper ion binding activity, role in cellular protein-containing complex assembly, copper ion transport and mitochondrial inner membrane, plasma membrane localization | CR_09300C_A/SCO1 |
| CPAR2_602990 | -9.79 | - Inf* | Ortholog(s) have copper uptake transmembrane transporter activity, role in copper ion import, high-affinity iron ion transport and cytoplasm, nucleus, plasma membrane localization | C6_00790C_A/CTR1 |

\*-Inf, no reads detected

Interestingly, CPAR2_602990, an orthologue of *C. albicans CTR1* (orf19.3646) with copper importing activity was the second most downregulated gene at 30 min (log_2_FC = −9.8) and no reads for this gene were detected after 60 minutes in our RNA sequencing data (Table 1, “-Inf”). Four other genes annotated as Cu transporters were also repressed at both time points.

The differential expression of these genes was validated by quantitative real-time PCR and we further tested for whether they would also respond to phagocytosis by primary human monocyte-derived human primary macrophages (MDMs). Only for the Cp*CRP1* gene was expression in response to both, *P. aurantium* and MDMs, in accordance, while the expression of Cp*CTR1* was regulated in an opposite manner when the yeast encountered MDMs (Fig. 4A+B). We further investigated the expression of Cp*CRP1* and Cp*CTR1* during *in vitro* Cu excess and depletion. As expected, the putative Cuexporter gene Cp*CRP1* showed induction when *Candida* was treated with 100 µM of Cu, and repression in the presence of the Cu chelator BCS (Fig. 4C). Even though the expression of Cp*CTR1* was not significantly influenced by the presence of BCS in the media, a remarkable downregulation of this gene was measured at high Cu concentration. It is noteworthy that differences in the expression levels for both genes, Cp*CRP1* and Cp*CTR1*, were more pronounced during encounters with *P. aurantium* than with macrophages or the metal itself.

**Fig. 4:**
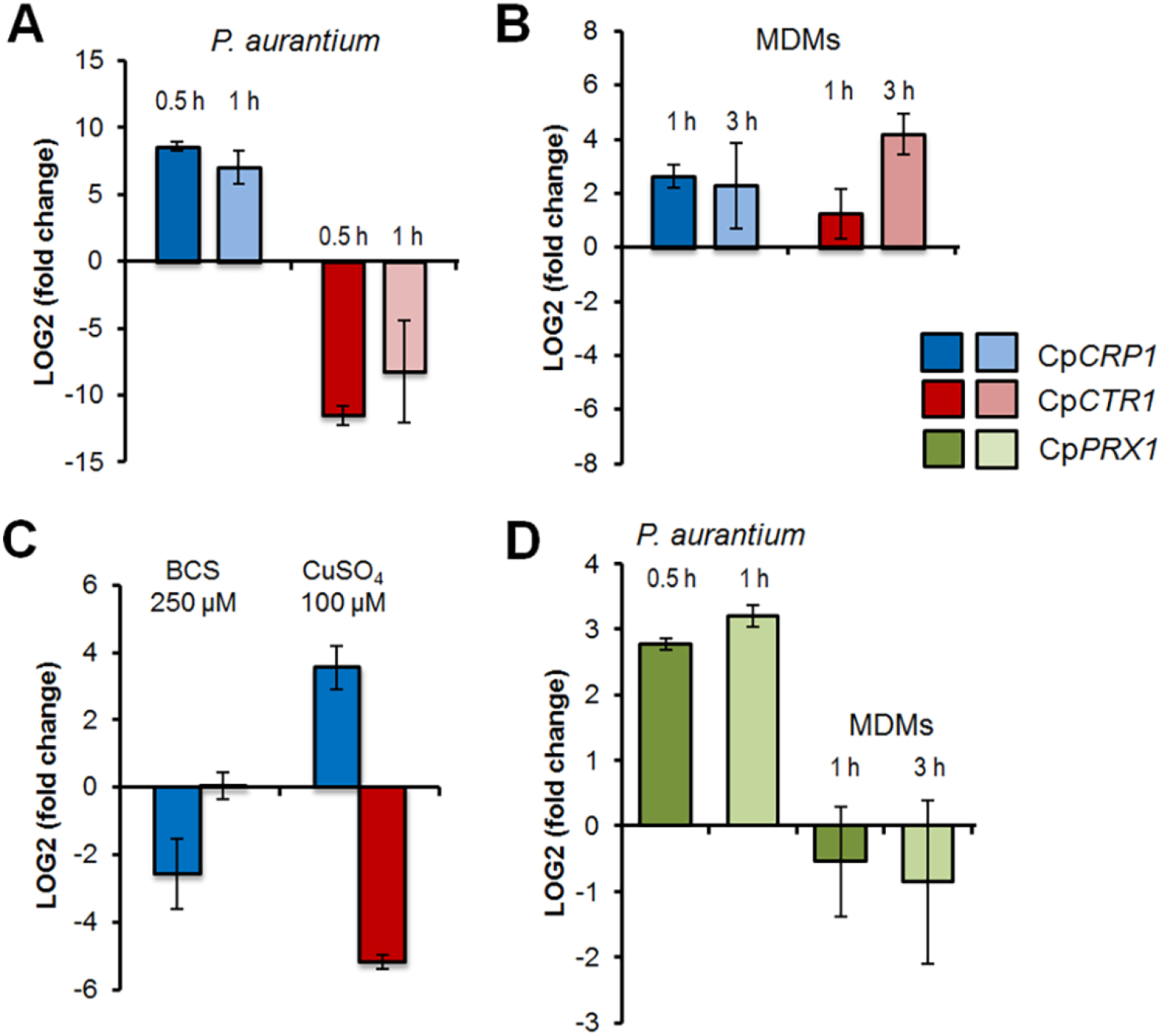
Expression of copper and redox homeostasis genes. Expression of the *CRP1* (CPAR2_203720), *CTR1* (CPAR2_602990), and *PRX1* (CPAR2_805590) of *C. parapsilosis* was analyzed by qRT-PCR using total RNA isolated after exposure to *P. aurantium* **(A, D)**, human monocyte-derived macrophages (MDMs, **B, D**), and in the presence of the copper ion chelator BCS or CuSO_4_ **(C)**. All data show average expression levels relative to time point 0 based on three biological and three technical replicates. Error bars indicate the standard deviation.

Intriguingly, four genes encoding superoxide dismutases (SODs) of the Cu/Zn type (*CPAR2_500330, CPAR2_500390, CPAR2_213540, CPAR2_213080*) and one Fe/Mn-SOD (*CPAR2_109280*) were strongly repressed in the presence of *P. aurantium* (Table 2). In contrast, genes involved in the thioredoxin antioxidant pathway were found to be highly upregulated, such as *CPAR2_304080*, *CPAR2_500130*, or *CPAR2_805590*. The latter being an orthologous gene to *C. albicans PRX1*, a thioredoxin-linked peroxidase shown to be primarily involved in the reduction of cellular organic peroxides (37). More than a 5-fold increase in transcript level was observed in *C. parapsilosis* after 30 min of co-incubation with *P. aurantium*. This upregulation further increased to 9-fold after another 30 min of co-incubation with the predator, however, remained unaffected in response to primary macrophages (Fig. 4D).

**Table 2:** Expression of *C. parapsilosis* genes* involved in response to oxidative stress (GO:0006979)

| name | log2FC<br>30 min | log2FC<br>60 min | description | *C. albicans* orthologue |
| --- | --- | --- | --- | --- |
| CPAR2_805590 | 2.03 | 2.9 | thioredoxin peroxidase activity and role in cell redox homeostasis, cellular response to oxidative stress, response to cadmium ion, sporocarp development involved in sexual reproduction | C7_02810W_A/PRX1 |
| CPAR2_101390 | 2.16 | 2.84 | NAD binding, nucleoside diphosphate kinase activity | C5_02890W_A/YNK1 |
| CPAR2_802460 | 3.24 | 3.75 | ribosomal large subunit binding, ribosomal small subunit binding activity and role in cellular response to oxidative stress, cytoplasmic translation | CR_00860C_A/TMA19 |
| CPAR2_213080 | -7.18 | -Inf** | Ortholog(s) have superoxide dismutase activity and role in cellular response to superoxide, evasion or tolerance by symbiont of host-produced reactive oxygen species | C2_00660C_A/SOD4 |
| CPAR2_803850 | -6.74 | -6.14 | Has domain(s) with predicted catalase activity, heme binding activity and role in oxidation-reduction process, response to oxidative stress | orf19.6229/CAT1 |
| CPAR2_806310 | -6.47 | -5.89 | ATPase activity, GTPase activity | C2_09220W_A/DDR48 |
| CPAR2_500330 | -5 | -6.92 | superoxide dismutase activity | orf19.2770.1/SOD1 |
| CPAR2_808660 | -4.59 | -4.53 | alditol:NADP+ 1-oxidoreductase activity and role in D-xylose catabolic process, arabinose catabolic process, cellular response to oxidative stress | C3_06860C_A |
| CPAR2_406810 | -4.42 | -4.59 | role in cellular response to oxidative stress, pathogenesis and plasma membrane localization | C2_06870C_A/PST1 |
| CPAR2_101680 | -3.97 | -5.25 | nitric oxide dioxygenase activity, nitric oxide reductase activity | CR_07790C_A/YHB1 |
| CPAR2_101350 | -3.67 | -2.87 | D-xylose:NADP reductase activity, NADPH binding, mRNA binding activity | C5_02930C_A/GRE3 |
| CPAR2_204330 | -3.52 | -3.07 | ADP binding, ATP binding, ATPase activity, coupled, chaperone binding, misfolded protein binding, unfolded protein binding activity | CR_08250C_A/HSP104 |
| CPAR2_806210 | -3.41 | -4.03 | role in NADH oxidation, positive regulation of apoptotic process, regulation of reactive oxygen species metabolic process, response to singlet oxygen and mitochondrion, nucleus localization | C2_08100W_A |
| CPAR2_804600 | -3.36 | -3.12 | Ortholog(s) have glutamate decarboxylase activity and role in cellular response to oxidative stress, glutamate catabolic process | C1_11660W_A/GAD1 |
| CPAR2_406510 | -3.35 | -4.7 | DNA-binding transcription factor activity, RNA polymerase II-specific, RNA polymerase II proximal promoter sequence-specific DNA binding activity | C2_07170C_A/AFT2 |
| CPAR2_208070 | -3.26 | -2.69 | superoxide dismutase copper chaperone activity and role in cellular copper ion homeostasis, cellular response to metal ion, protein maturation by copper ion transfer, removal of superoxide radicals | C1_07180W_A/CCS1 |
| CPAR2_103080 | -2.63 | -2.89 | glyoxalase III activity and role in cellular response to nutrient levels, cellular response to oxidative stress, methylglyoxal catabolic process to D-lactate via S-lactoyl-glutathione | C3_02610C_A/GLX3 |
| CPAR2_209930 | -2.54 | -3.54 | DNA-binding transcription factor activity, RNA polymerase II-specific activity | C2_05860C_A |
| CPAR2_109280 | -2.51 | -3.01 | manganese ion binding, superoxide dismutase activity | C1_01520C_A/SOD2 |
| CPAR2_808750 | -2.47 | -2.08 | alditol:NADP+ 1-oxidoreductase activity, glycerol dehydrogenase [NAD(P)+] activity, mRNA binding activity | C3_07340W_A/GCY1 |
| CPAR2_601800 | -2.4 | -1.84 | cytochrome-b5 reductase activity, acting on NAD(P)H activity and role in cellular response to oxidative stress, ergosterol biosynthetic process | C6_02040W_A/MCR1 |
| CPAR2_704130 | -2.39 | -2.77 | role in cellular response to drug, cellular response to oxidative stress and filamentous growth of a population of unicellular organisms in response to biotic stimulus, more | C7_03220C_A/ZCF29 |
| CPAR2_804070 | -2.34 | -3.4 | DNA-binding transcription factor activity, RNA polymerase II-specific, RNA polymerase II regulatory region sequence-specific DNA binding activity | CR_00630W_A |
| CPAR2_105790 | -2.28 | -2.5 | disulfide oxidoreductase activity, glutathione peroxidase activity, glutathione transferase activity and role in cellular response to oxidative stress, glutathione metabolic process, pathogenesis | C1_00490C_A/TTR1 |
| CPAR2_200560 | -2.04 | -2.15 | S-(hydroxymethyl)glutathione dehydrogenase activity, alcohol dehydrogenase (NAD) activity, hydroxymethylfurfural reductase (NADH) activity | CR_10250C_A/FDH3 |
| CPAR2_301730 | -1.88 | -2.27 | DNA-binding transcription factor activity, RNA polymerase II-specific, proximal promoter sequence-specific DNA binding activity | C1_08940C_A/MSN4 |
| CPAR2_500390 | -1.75 | -1.73 | superoxide dismutase activity | C4_02320C_A/SOD1 |
| CPAR2_204160 | -1.72 | -2.44 | role in cellular response to oxidative stress, chromatin silencing at silent mating-type cassette, pathogenesis | CR_05390W_A/PST3 |
| CPAR2_600070 | -1.59 | -1.73 | actin filament binding, protein binding, bridging activity | C6_02730W_A/SAC6 |
| CPAR2_102830 | -1.37 | -1.72 | Ortholog(s) have cytochrome-c peroxidase activity, role in cellular response to reactive oxygen species and mitochondrial intermembrane space, mitochondrial membrane localization | C3_02480C_A/CCP1 |

\*only genes with p <0.01 according to EdgeR are displayed
\*\*.-Inf, no reads detected

### *C. parapsilosis* is exposed to ROS during phagocytosis by *P. aurantium*

The oxidative burst leading to production of ROS occurs frequently when phagocytes encounter microbial prey. The repression of catalases and superoxide dismutases, but concurrent induction of genes involved in redox homeostasis prompted us to analyze whether *C. parapsilosis* is exposed to ROS during interaction with the fungivorous predator. When co-incubating *C. parapsilosis* with *P. aurantium* in the presence of the superoxide (∙O_2_^−^) indicator dihydroethidium (DHE), an increase in red fluorescence of cultures was specific to the presence of amoebae and reached a maximum after 10 min of co-incubation (Fig. 5A). Fluorescence microscopy of single cells of *P. aurantium* using either DHE or the alternative ROS sensor CellROX^®^Deep Red further revealed that ROS production was locally specific to *P. aurantium* actively feeding on *C. parapsilosis* (Fig. 5B), suggesting that yeast cells are exposed to increased levels of ROS upon phagocytic processing in *P. aurantium*.

**Fig. 5:**
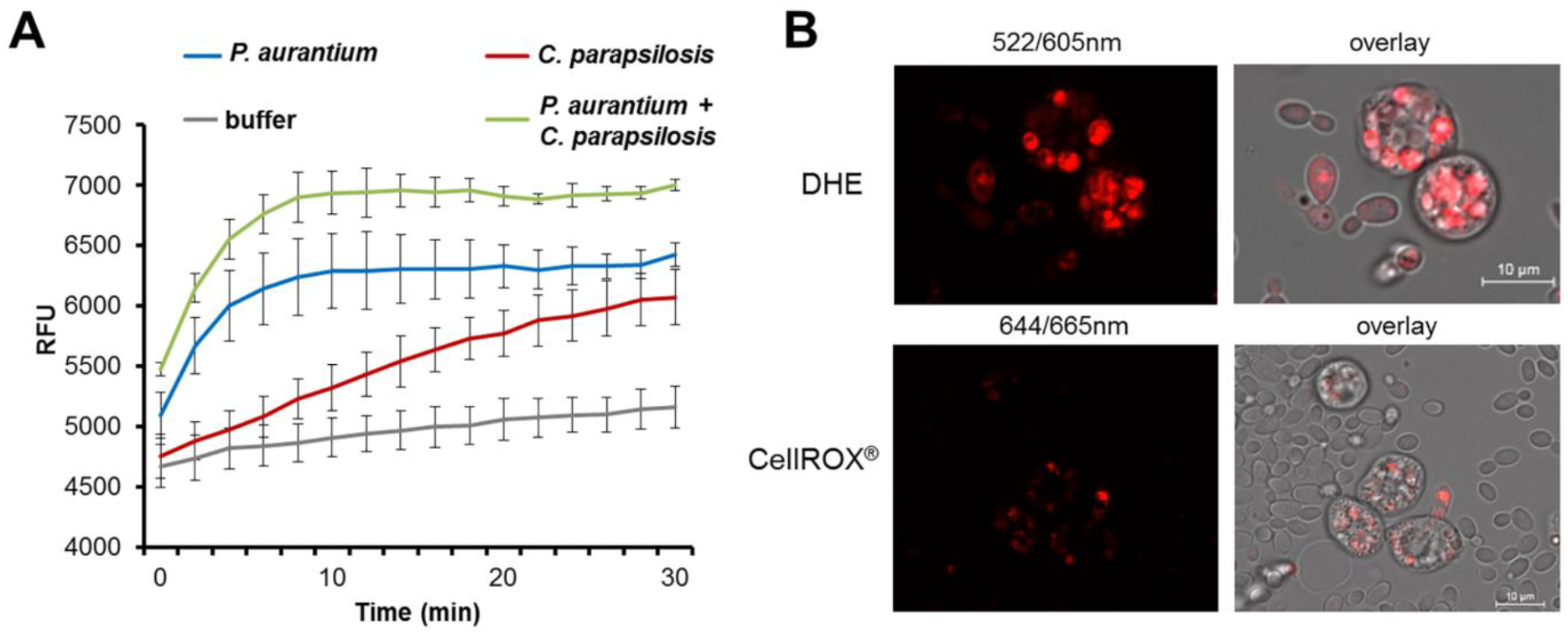
ROS production by *P. aurantium* during phagocytosis of *C. parapsilosis.* **(A)** ROS were determined indirectly as the increase in DHE oxidation over 30 min in co-incubations of *C. parapsilosis* with *P. aurantium.* Data represent mean RFU (λ_ex_ 522/ λ_em_ 605 nm) of three independent samples over 30 min. **(B)** ROS production was primarily localized to feeding cells of *P. aurantium.* Cells were co-incubated with *C. parapsilosis* in the presence of ROS sensitive probes DHE or CellROX^®^ Deep Red and images were taken after 30 min.

### Copper and redox homeostasis contribute to the resistance against *P. aurantium* and macrophages

The expression profile and its similarity to its orthologue in *C. albicans* suggested a role for Crp1 of *C. parapsilosis* in detoxification of high Cu levels. Deleting Cp*CRP1* (*∆/∆crp1*) displayed no apparent growth defect in SD medium at 30°C and the mutant strain tolerated even high concentrations of Cu above 1 mM. Its sensitivity towards this transition metal changed dramatically when cells were exposed to a more acidic pH on solid or in liquid media (Fig. 6). At a pH of 3, Cp*CRP1* proved to be important for growth at Cu concentrations between 500 and 1000 µM, indicating that the function of Crp1p could be crucial under the acidic conditions of the phagolysosome.

**Fig. 6:**
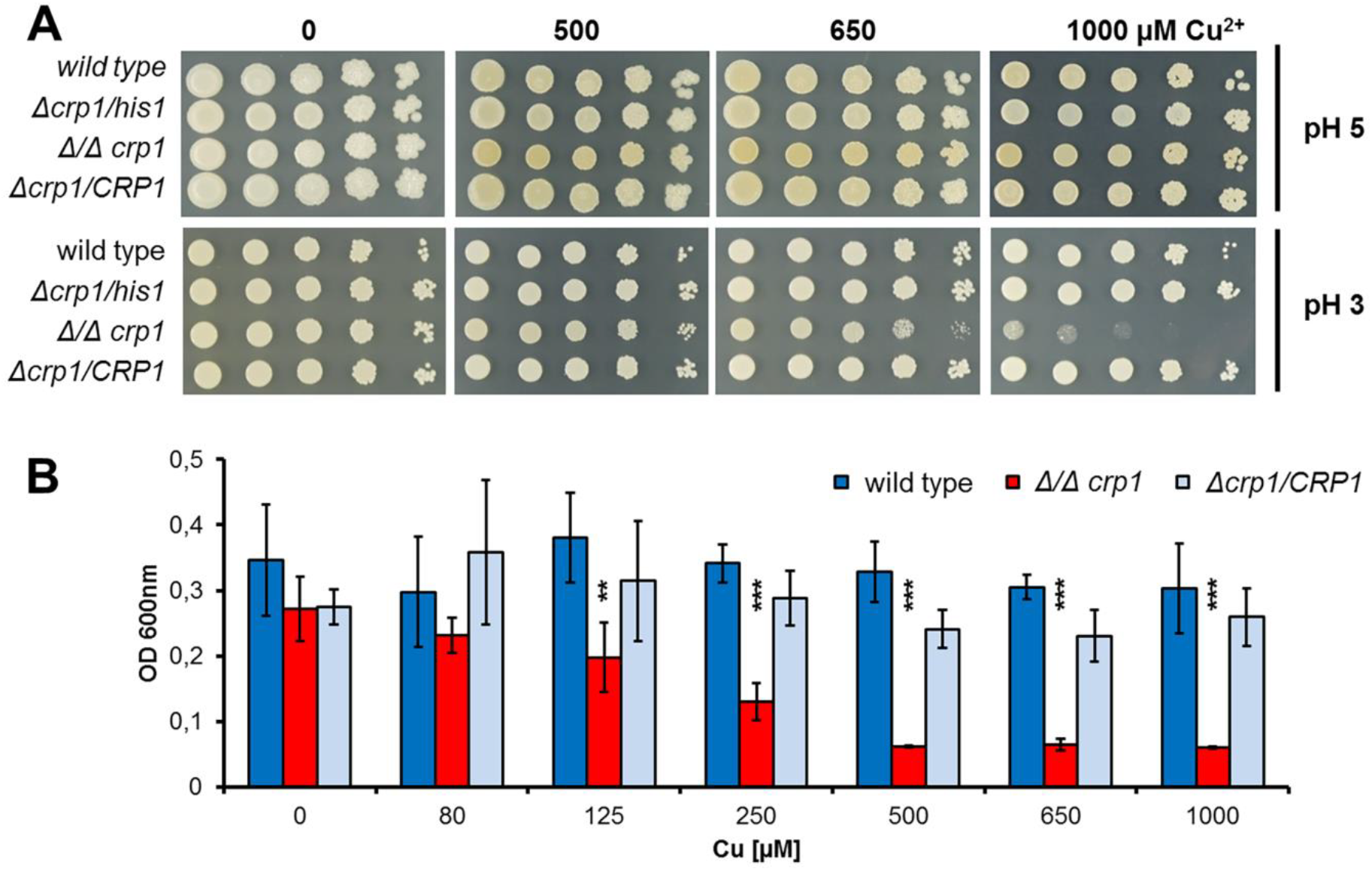
Crp1p protects *C. parapsilosis* from high Cu levels at acidic pH. **A.** The *Δ/Δcrp1* mutant strain showed a pH-dependent copper sensitivity in comparison to the wild type during growth at pH 3. **B.** Increased sensitivity of the *Δ/Δcrp1* mutant strain to high Cu concentrations in liquid malt extract (pH 3) compared to the wild type and complemented strain. Data represent the mean and standard deviation of three biological replicates with asterisks indicating statistical significance in an unpaired Student’s t-test between the values obtained for the *Δ/Δcrp1* strain and the wild type (∗∗∗, p < 0.001).

We also addressed the antioxidant function of *PRX1* in *C. parapsilosis*, by subjecting a homozygous mutant (*Δ/Δprx1*) to oxidative stress delivered by hydrogen peroxide (H_2_O_2_) and *tert*-butyl hydroperoxide (*t*-bOOH). The sensitivity of the mutant towards H_2_O_2_ was nearly indistinguishable from the wild type and the organic peroxide had only a mild effect on the growth of *Δ/Δprx1* on solid medium supplemented with adenine, uracil, and 9 amino acids (Fig. 7A). However, the impact of oxidative stress was more severe when these supplements were omitted from the medium. Under these conditions, growth in liquid medium was significantly reduced for *Δ/Δprx1* even in the absence of an external stressor (Fig. 7B). When using 11 selective dropouts, each one lacking a singlecomponent, we found that a lack of methionine was responsible for the growth defect of *Δ/Δprx1.* The omission of methionine from the normal medium, in combination with the organic peroxide affected the wild type and the mutant strains to similar extents (Fig. 7C).

**Fig. 7:**
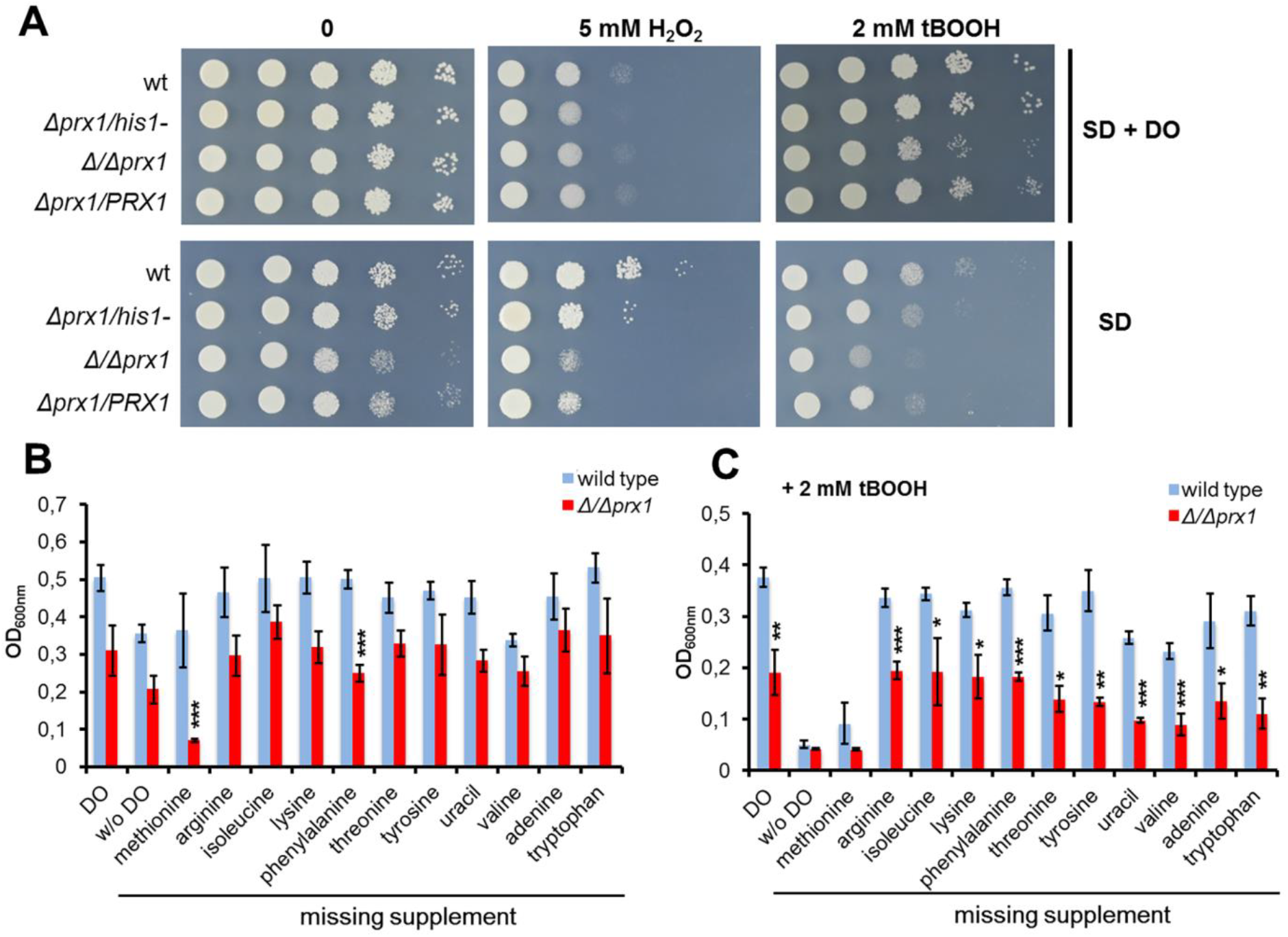
The antioxidant role of *PRX1* in *C. parapsilosis*. **A.** Growth of *C. parapsilosis* on solid SD media with or without drop-out supplement (DO) in the presence of tBOOH or H_2_O_2_ as oxidative stressors. **B and C.** Growth of the wild type (wt) and the *Δ/Δprx1* mutant in liquid SD media, supplemented with drop-out solutions, selectively missing one essential component (B) and in the presence of tbOOH (C). Growth was measured as optical density at 600 nm. Data represent the mean and standard deviation of three biological replicates with asterisks indicating statistical significance in an unpaired Student’s t-test between the values obtained for the *Δ/Δprx1* mutant in comparison to wt (∗p < 0.05, ∗∗ p < 0.01, ∗∗∗p < 0.001).

Both *CRP1* and *PRX1* are widely conserved across the *Candida* clade, including several species without any record as commensals or pathogens (Fig. S3). To test whether these two genes contribute to the defense against an environmental predator, the deletion mutants for *CRP1* and *PRX1* were confronted with *P. aurantium.* Both mutants showed decreased survival in comparison to the wild type after 3 hours of co-incubation (Fig. 8A). As we hypothesized that both mechanisms for stress defence could also contribute to survival when encountering human innate immune cells, we performed another co-incubation assay with primary macrophages. Both *Δ/Δcrp1* and *Δ/Δprx1* displayed reduced survival when confronted with MDMs (Fig. 8B), indicating that these genes not only mediate resistance to copper and oxidative stress during predation by amoeba but could also play a role during immune evasion in a human host.

**Fig. 8:**
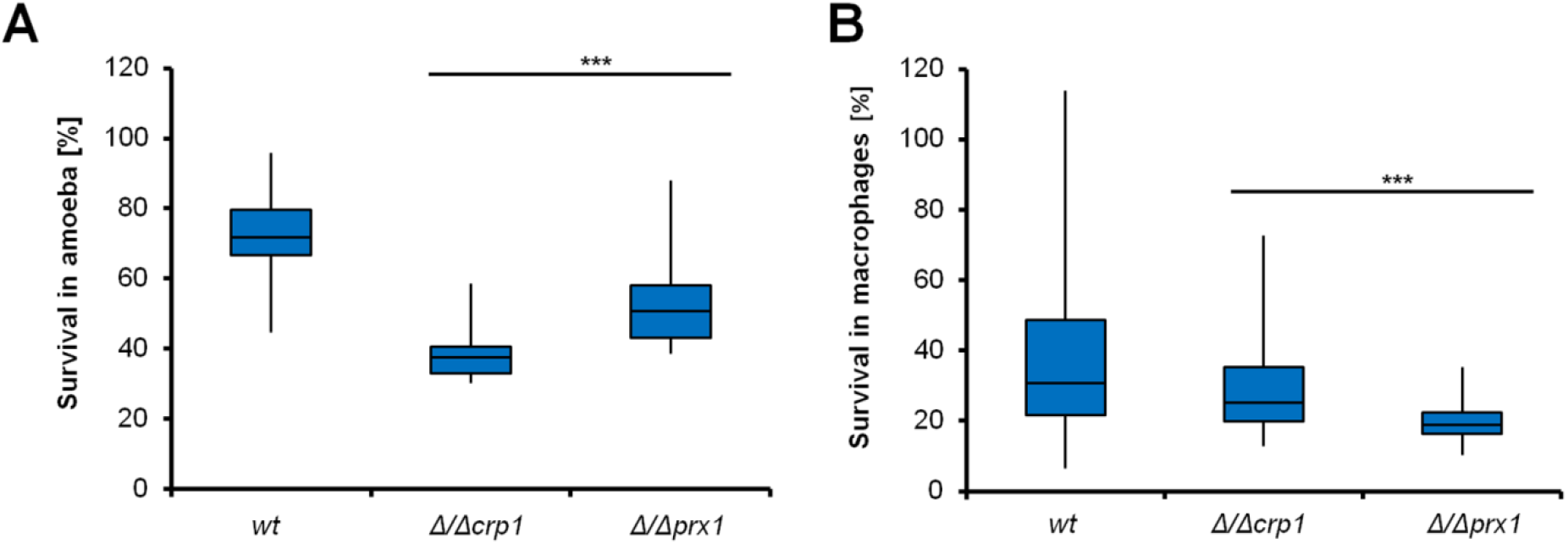
Survival of *C. parapsilosis* strains during amoeba predation (A) and phagocytosis by primary macrophages (B). Strains of *C. parapsilosis* were incubated with *P. aurantium* for 3 h at a yeast-to-amoeba ratio of 10:1 **(A)** or with primary macrophages isolated from at least 6 different anonymous donors at a yeast-to-macrophage ratio of 1:1 **(B)**. The number of survivors was determined by plating the cells on YPD media and counting the CFUs. The boxes signify the 25^th^ and 75^th^ percentile. The median is represented by a short black line within the box for each strain. The whiskers indicate the highest and lowest values from three independent biological and six technical replicates. Asterisks show statistical significance in an unpaired Student’s t-test between the values obtained for the null mutants in comparison to the parental strain, wt (∗∗∗, p<0.001).

## Discussion

The arms race between phagocytic predators and their microbial prey is thought to have shaped virulence determinants of bacteria and fungi (21, 38). Amoebae are predominant environmental micro-predators, but only a few of them have been described to actively feed on fungi (39–41). Such a fungivorous lifestyle has been described for *Protostelium mycophagum*, the type species for the polyphyletic group of protosteloid amoebae that form microscopic, stalked fruiting bodies from single cells and are found on nearly all continents (42–45). We have recently isolated and characterized a strain of *P. aurantium* (formerly known as *Planoprotostelium aurantium*), which was found to selectively recognize, kill, and feed on a wide range of ascomycete and basidiomycete yeasts, including major human pathogens of the *Candida* clade (32). *C. parapsilosis* acted as a preferred food source, while *C. albicans* was found to be protected from initial recognition by an extensive coat of mannoproteins, and *C. glabrata* showed delayed processing after ingestion. A similar survival strategy seems to rescue *C. glabrata* when encountering macrophages. Here, its ability to persist and even replicate inside the phagocyte has been well documented and characterized to the level of single genes (15, 16, 46). A functional genomic approach identified 23 genes in *C. glabrata* which were critically involved in the survival of macrophage phagocytosis (47). When comparing this set of 23 genes to all genes expressed during predation by *P. aurantium*, we found 7 genes to be highly upregulated (log_2_FC >1.5) at both time points. The three most upregulated genes with a log_2_FC of more than 2 were *CgGNT1* (CAGL0I09922g), *CgOST6* (CAGL0G07040g), and *CgPMT2* (CAGL0J08734g); all involved either in cell wall modification or protein glycosylation.

All these genes share orthologues with the other two *Candida* species, but when confronted with *P. aurantium*, only *PMT2* was upregulated in *C. parapsilosis* and even more so in *C. albicans*. In the latter, the gene encodes an essential protein, O-mannosyltransferase, which renders the cell more resistant to antifungals and cell wall perturbing agents (48, 49). The upregulation of mannan synthesis in *C. albicans* in the presence of the predator seems not to be limited to O-linked mannans, but was also observed for the N-linked type. *MNN2* and *MNN22* are two members of another well-characterized family of N-mannosyltranferses whose absence severely affects the mannoprotein coat of *C. albicans* (50, 51). Both genes showed induction levels comparable to *PMT2*. The pivotal role of the mannan coat of *C. albicans* during an interaction with phagocytes of the innate immunity is well studied, as defective O-and N-linked mannosylation led to an increased uptake and phagosomal maturation, most likely through unmasking of β-glucans and enhanced recognition of *C. albicans via* the Dectin-1 receptor (52, 53). The fact that mannan biosynthesis was upregulated in *C. albicans* is in agreement with the previous finding that mannosidase-treated cells were internalized more frequently by *P. aurantium* (32).

The different interaction patterns of the three yeasts were partially reflected throughout the transcriptome of their orthologous genes. Although, a large gene set was induced in *C. glabrata*, this showed relatively low variation over time, indicating that *C. glabrata* responds to the presence of the amoeba, but not to predation or killing. This may explain why the general response to the predator comprised only 79 orthologues. Of these, 48 were commonly induced, among them the orthologues of *CDC54*, *CDC46*, and *MCM3*, indicating that all three yeast species were metabolically active in the M/G1 phase of the cell cycle (54). Genes involved in fatty acid catabolism were generally induced while their biosynthesis was rather repressed. In contrast, amino acid biosynthesis was commonly upregulated, and as this was seen also for the non-internalized *C. albicans*, it presumably results rather from the response to the nutrient-deprived growth medium used during the confrontation than from direct interaction with the amoeba.

Of over 1,500 orthologous genes that were differentially regulated in either *C. albicans* or *C. parapsilosis*, only 251 were common DEOs for both species. Within this gene set, the impact on Cu homeostatic genes was preeminent, especially for *C. parapsilosis*, and the null mutant for *CRP1*, the gene with the highest induction, which pointed towards a vital role of copper during predation by *P. aurantium*. Intoxication by copper is especially effective in highly acidic environments as occur during early maturation of the phagolysosome and has been shown as an effective strategy of macrophages to control the primary intracellular pathogen *Mycobacterium tuberculosis* (55). Of all three species, inhibition of phagolyosomal acidification has exclusively been reported for *C. glabrata* (16), which might explain why Cu resistance genes were not found to not respond in this yeast. Elevating copper and ROS within its acidic phagolysosome was also found for the bacteriovorous amoeba *Dictyostelium discoideum* and has most likely contributed to the spread of copper resistance islands among bacterial pathogens (56). From this perspective, it cannot be surprising that highly tuned copper homeostatic systems were elucidated in the major environmentally acquired fungal pathogens *Aspergillus fumigatus* and *Cryptococcus neoformans* (57, 58). Both fungi also exploit similar escape strategies when confronted with amoebae or mammalian phagocytes (23, 25, 59). A recent screening approach identified Sur7 as a Cu-protective protein which reduces membrane permeability to Cu in *C. albicans* (60). Although downregulation of *CaSUR7* was observed at both time points after confrontation with the amoeba, its putative *C. parapsilosis* orthologue (CPAR2_602600) showed higher expression after one hour of co-incubation with *P. aurantium*. Also, for *C. glabrata* which seems to lack a *CRP1* orthologue, *CgSUR7* (CAGL0L01551g) was highly induced at both time points. At least some of the toxic effects of Cu could well be inflicted via Fenton-type chemistry with ROS, which were actively produced in feeding *P. aurantium.* Their impact on the yeast would likely be further aggravated by the downregulation of nearly all SODs, as seen for *C. parapsilosis*, and, to a lesser degree, also for *C. albicans* during confrontation with the predator. At least for *C. albicans* it is well known that it can adapt the expression of its SOD genes according to the availability of the metal cofactors (61), thus high levels of copper and ROS, as seen here, should activate the expression of genes encoding the SOD of the Cu/Zn type. However, these genes in particular are severely repressed in *C. albicans* and *C. parapsilosis* (Table S3), suggesting that the predator could interfere with the normal response to ROS via a yet unknown mechanism.

The one-cysteine peroxiredoxin *PRX1* was among the very few oxidative stress genes which expression was increased in *C. parapsilosis*. Its orthologous gene was also upregulated during in *C. albicans* during co-incubation with macrophages (62). We found that an essential cellular function of *PRX1* is tightly linked to a lack of methionine. Intriguingly, among the only 48 commonly upregulated genes in all three *Candida* species, 5 are involved in the metabolism of sulfur-containing amino acids (*ECM17, MET15, MET16, MET3, SAM2*). For all five genes, the induction was lowest for *C. glabrata*. Amino acid deprivation, and more specifically a limitation in methionine, also occurs in the phagolysosome of neutrophilic granulocytes (63, 64). Sensitivity to ROS is phenotypically well established for the methionine biosynthesis pathway in baker’s yeast, as mutants lacking either SOD1 or its chaperone CCS1 were unable to grow in normoxic environments due to a methionine auxotrophic phenotype (65, 66).

In conclusion, our results indicate that the fungivorous feeding by the predator *P. aurantium* activates the fungal Cu and redox homeostasis which is essential to shield methionine biosynthesis from oxidative inactivation. After millions of years of coevolution of amoebae and fungi, it is well conceivable that such basic molecular tools for resistance against environmental phagocytes proved to be valuable for survival in phagocytic cells of higher eukaryotes, like humans and mammals.

## Material and Methods

### Strains and growth conditions

*Protostelium aurantium var. fungivorum* has been isolated in Jena, Germany, as described previously (31). Isolated amoebae were further grown in standard-size Petri dishes (94×16 mm, Greiner Bio-One, Austria) in PBS (80 g l^−1^ NaCl, 2 g l^−1^ KCl, 26.8 g l^−1^ Na_2_HPO_4_ x 7 H_2_O, 2.4 g l^−1^ KH_2_PO_4_, pH 6.6) with *Rhodotorula mucilaginosa* as a food source at 22°C, if not stated differently. All yeast strains are listed in Table S4. If not indicated otherwise, all fungi were grown in YPD medium (1 % yeast extract, 2 % peptone, 2 % glucose) at 30°C, supplemented with 1.5 % [w/v] agar for growth on solid media. Mutant strains of *C. parapsilosis* were grown on SD-agar (0.4 % [w/v] yeast nitrogen base with ammonium sulfate, 2 % [w/v] glucose, 1.5 % [w/v] agar), supplemented with 10 % [v/v] of a selective drop-out solution excluding leucine or histidine. Complemented strains were grown on YPD agar supplemented with 100 µg ml^−1^ nourseothricin (clonNAT, Werner BioAgents, Jena, Germany).

### Confrontation of *Candida sp.* with *P. aurantium*

*P. aurantium* was pre-cultured in 20 x wMY medium at 22°C and cells were washed with fresh medium, scraped from the surface, harvested by centrifugation for 10 min at 800 x g and resuspended in 20 x wMY (40 mg l^−1^ yeast extract, 40 mg l^−1^ malt extract, 0.75 g l^−1^ K_2_HPO_4_). *Candida sp.* were grown overnight in YPD medium at 30°C, harvested, washed twice with cold 20 x wMY. Yeast cells were resuspended and adapted in fresh 20 x wMY at room temperature. Amoebae and yeast cells were counted in an automatic cell counter (Casy^®^TT Cell Counter, OLS Bio, Germany). Confrontations were carried out by spreading mixtures of both interaction partners at prey-predator ratios of 10:1 on 20 x wMY agar-plates (120 x 120 x 17 mm, Greiner Bio-One). At indicated time points cells were washed off the plate and centrifuged at 3000 rpm, 5 min. Pellets were immediately frozen in liquid N_2_ and used for the isolation of total RNA. Four independent amoeba and yeast cultures were used in this experiment.

### RNA isolations from yeast cells and co-cultures with *P. aurantium*

Frozen cell pellets from yeast cultures or co-incubations with *P. aurantium* were resuspended in TES buffer (10 mM Tris-HCl, pH 7.5, 10 mM EDTA, 0.5% [w/v] SDS) and transferred into a chilled tube containing zirconia beads (ZYMO Research, Irvine, CA, USA). Primary extractions of RNA were performed with acidic phenol:chloroform (5:1) shaking at 1,500 rpm at 65°C for 30 min in a thermoblock. Afterwards, samples were frozen at −80°C for 30 min, centrifuged at 10,000 g for 15 min for phase separation. Samples underwent two more extractions using phenol:chloroform (5:1) and chloroform:isoamyl alcohol (24:1). Total RNA was equilibrated with 10% [w/v] of 3 M sodium acetate (pH 5.2) and precipitated in ice-cold ethanol. After centrifugation, precipitates were washed in 70 % [v/v] ethanol, air dried, and dissolved in nuclease-free water. RNA samples were stored at −80°C until use.

### RNA isolations from yeast cells after confrontation with monocyte-derived macrophages (MDMs)

Adherent macrophages with attached and ingested yeast cells were washed and subsequently lysed by adding RLT lysis buffer containing β-mercaptoethanol (Qiagen, Hilden, Germany) and shock-freezing the plate in liquid nitrogen. Cells were detached by scraping and transferred into screw cap tubes, sedimented by centrifugation, and washed once with RLT buffer to remove most of the host RNA. Yeast pellets were shock-frozen again in liquid nitrogen and stored at −80°C. For RNA isolation, pellets were resuspended in 400 µl of AE buffer (50 mM sodium acetate pH 5.3 and 10 mM EDTA) and 40 µl of 10 % [w/v] SDS. After mixing for 30 sec, cells were extracted with phenol: chloroform: isoamyl alcohol [25:24:1] for 5 min at 65°C and then frozen at −80°C. Phase separations, precipitation, and resuspension of the purified RNA were performed as above.

### RNA sequencing and analysis of expression data

The preparation of cDNA libraries from total RNA and the sequencing was performed at LGC Genomics GmbH (Berlin, Germany). Briefly, the quality of RNA samples was first controlled using a 2100 Bioanalyzer (Agilent, CA, USA). Next, samples were enriched for mRNA using oligo-dT binding and magnetic separation using the NEBNext Poly(A) Magnetic Isolation Module (New England Biolabs). Samples were reverse transcribed using the NEBNext RNA First and Second Strand Synthesis Modules (New England Biolabs) and purified. The Encore Rapid DR Multiplex system (Nugen) was used for preparation of cDNA-libraries which were amplified in a volume of 100 µl for 15 cycles using MyTaq (Bioline) and standard Illumina primers. From these libraries, 2 x 75 bp (*C. parapsilosis*) or 2 x 150 bp (*C. albicans* and *C. glabrata*) paired-end reads were sequenced on an Illumina MiSeq platform. FastQC (67) and Trimmomatic v0.32 (68) were used for quality control and trimming of library adaptors. Mapping of reads was achieved with TopHat2 v2.1.0 (69) against the reference genomes of *C. parapsilosis*, *C. glabrata*, and *C. albicans* in combination with the genome of *P. aurantium*. Differential gene expression between time points was analyzed with EdgeR (70). A list of all differentially expressed genes is provided as Dataset S1. All sequencing data is available from the GEO repository under the accession number GSE116535. The Venn diagram was computed using the package “VennDiagram” from the statistical programming language R. The genes in all areas of the Venn diagram are listed in Dataset S4. The PCA was conducted using the method “prcomp” from the “stats” package of R.

### Gene ontology analysis

Gene ontology (GO) clustering analysis was performed on all differentially up- and downregulated genes of three *Candida* species using GO Slim (71) - Mapper tool available at the Candida Genome Database (http://www.candidagenome.org/cgi-bin/GO/goTermMapper) for biological process, molecular function, and cellular component. All data from the Gene Ontology analysis is provided as Dataset S2.

### Quantitative real-time reverse transcription-PCR (qRT-PCR)

For all qRT-PCR reactions 1µg of total RNA was treated with DNase using RQ1 RNase-free DNase (Promega, USA) and transcribed into cDNA (RevertAid First Strand cDNA Synthesis Kit, Thermo Scientific). The cDNA was diluted 1:10 and used for qRT-PCR with SYBR Select Master Mix (Applied Biosystems) performed in a thermocycler (Step One Plus, Applied Biosystems). The experiments were done in three biological and three technical replicates. The expression rates reported here are relative to the expression values of the housekeeping gene *ACT1* of *C. parapsilosis*. All primers are listed in Table S5.

### Detection of reactive oxygen species

The production of reactive oxygen species (ROS) during amoeba predation was measured using dihydroethidium (DHE; Thermo Fisher, Dreieich, Germany) at a final concentration of 10 µM. Amoebae and yeasts were seeded at an MOI of 10. Increased fluorescence, indicating ROS production, was measured using an Infinite M200 Pro fluorescence plate reader (Tecan, Männedorf, Switzerland) in intervals of 2 min over a 30 min period at λ_ex_ 522 nm/λ_em_ 605 nm. ROS production was further visualized by using DHE staining as mentioned above or CellROX^®^Deep Red staining (Thermo Fisher) at a final concentration of 5 µM. Fluorescence images were captured using the Zeiss Axio Observer 7 Spinning Disk Confocal Microscope (Zeiss, Germany) at the λ_ex_ 370 nm/λ_em_ 420 nm (for non-oxidized DHE), λ_ex_ 535 nm/λ_em_ 610 nm (for oxidized DHE), and at λ_ex_ 640 nm/λ_em_ 665 nm for CellROX^®^Deep Red.

### Construction of gene deletions and complementations in *C. parapsilosis*

Target genes were deleted from the leucine and histidine auxotrophic parental strain CLIB2014 using a fusion PCR method described previously (72) and adapted to *C. parapsilosis* (73). All primer sequences and target genes are listed in Table S5. Briefly, approximately 500 bp of the upstream and downstream DNA loci of the coding sequence were amplified by PCR with the primer pairs P1/P3 and P4/P6, respectively. The selectable markers, *C. dubliniensis HIS1* and *C. maltosa LEU2* genes were amplified with the P2/P5 primer pair from the plasmids pSN52 and pSN40, respectively. All PCR products were further purified using Gel/PCR DNA Fragments Extraction Kit (Geneaid Biotech, New Taipei City, Taiwan) and connected *via* PCR through overlapping sequences of the P2/P3 and P4/P5 primer pairs. The entire deletion cassette was amplified using primers P1 and P6 and transformed into the recipient strain in two rounds of transformation. The first allele was replaced by the *CmLEU2* marker and the second allele with the *CdHIS1* marker. Site-specific integration of the selection marker was checked by PCR at both ends of the deletion constructs. Loss of expression was also confirmed by qRT-PCR targeting the respective ORF (Fig. S4A-D).

To generate complemented strains, the neutral locus NEUT5L was targeted as described in (74). The promoter-OR-terminator regions were amplified from the CLIB214 parental strain using specific rec_F/R primers listed in Table S5. The dominant nourseothricin resistance marker *NAT1*, and a modified sequence of *C. parapsilosis* NEUT5L locus, were amplified from plasmid pDEST_TDH3_NAT_CpNEUT5L_NheI using Clon_F/Clon_R primers. The 5’ tails of *gene name_*rec_F/*gene name_*rec_R primers contained flanking regions complementary to the sequence of Clon_F/Clon_R primers to allow fusion *via* circular polymerase extension cloning (CPEC cloning). The completely assembled plasmids (Fig. S4E) were directly used for transformation of *E. coli* DH5α without further purification. After purification from *E. coli* up to 3 µg of plasmid were enzymatically digested with *Stu*I or *Hpa*I and *Eco*RI to confirm their correct size. The modified sequence of *C. parapsilosis* NEUT5L locus contains a specific restriction site for *Stu*I which linearised the plasmid and enables the integration of the vector into the NEUT5L locus of *C. parapsilosis via* duplication. Integration of the vector into the genome of the parental strain was confirmed by PCR.

### Chemical transformation of *C. parapsilosis*

Overnight cultures of *C. parapsilosis leu2*Δ/*his1*Δ were diluted to an OD_600_ of 0.2 in YPD media and grown at 30°C/180 rpm to an OD_600_ of 1. The culture was harvested by centrifugation at 4,000 g for 5 min and the pellet was suspended in 3 ml of ice-cold water. After collecting the cells, the pellet was resuspended in 1 ml of TE with LiAc (0.1 M lithium acetate, 10 mM Tris-HCl, 1 mM EDTA, pH 7.5), followed by centrifugation for 30 s at 14,000 g. Cells were then suspended in 200 µl of the ice-cold TE-LiAc buffer. For transformation, 10 µl of boiled herring sperm DNA (2 mg/ml) and 20 µl of transforming DNA were added to 100 µl of competent cells. The mixture was incubated at 30°C without shaking for 30 min followed by the addition of 700 µl of PLATE solution (0.1 M lithium acetate, 10 mM Tris-HCl, 1 mM EDTA, pH 7.5 and 40 % PEG 3350). Afterwards, the samples were incubated overnight at 30 °C. The next day, samples were heat-shocked at 44 °C for 15 min, centrifuged, and washed twice with YPD medium. Following incubation in 100 µl of YPD for 2 hours (180 rpm, 30°C), samples were plated on SD agar plates, supplemented with essential amino acids, and either histidine or leucine to obtain heterozygous mutant strains. To select for homozygous mutant, histidine and leucine were omitted from the medium. Selective plates were incubated for two days at 30°C. To select for complemented strains, cells were plated on YPD agar with 100 µg ml^−1^ of nourseothricin.

### Sensitivity assays

Yeasts were grown overnight in YPD at 30°C/180 rpm, harvested by centrifugation at 10,000 g for 1 min, and washed twice with PBS. For droplet assays, cells were diluted to the concentration of 5×10^7^ ml^−1^ and 5 µl of serial 10-fold dilutions were dropped on agar plates. To determine the MIC_50_ of Cu, 2.5×10^4^ cells were seeded in a 96 well plate with malt extract broth buffered to pH 3 and CuSO_4_. For oxidative stress sensitivity assays, 11 selective drop-out solutions, each missing one component, were added to liquid SD medium (0.4 % [w/v] yeast nitrogen base with ammonium sulfate, 2 % [w/v] glucose) with or without 2 mM of *t*-BOOH (Luperox^®^ TBH70X, Sigma-Aldrich, USA) and approx. 3×10^2^ cells were seeded in 48 well plates. All plates were incubated at 30°C for two days. Growth in well plates was evaluated by measuring the optical density (OD_600_) in a plate reader (Infinite M200 Pro, Tecan, Männedorf, Switzerland). Data represent the average of 3 biological replicates.

### Isolation of monocyte-derive macrophages (MDMs)

Human peripheral blood mononuclear cells (PBMCs) were isolated by density centrifugation. PBMCs from buffy coats donated by healthy volunteers were separated through Lymphocytes Separation Media (Capricorn Scientific, Germany) in Leucosep™ centrifuge tubes (Greiner Bio-One). Magnetically labelled CD14 positive monocytes were selected by automated cell sorting (autoMACs; Miltenyi Biotec, Germany). To differentiate monocytes into MDMs, 1.7×10^7^ cells were seeded into 175 cm² cell culture flasks in RPMI 1640 media with L-glutamine (Thermo Fisher Scientific) containing 10 % heat-inactivated fetal bovine serum (FBS; Bio&SELL, Germany) and 50 ng ml^−1^ recombinant human M-CSF (ImmunoTools, Friesoythe, Germany) and incubated for five days at 37 °C and 5 % CO_2_ until the medium was exchanged. Stimulation with human M-CSF favours the differentiation to M2-type macrophages. After two additional days, adherent MDMs were detached with 50 mM EDTA in PBS and seeded in 6-well plates (for expression analysis) or in 96-well plates (for killing assay) to a final concentration of 1×10^6^ or 4×10^4^ MDMs/well, respectively in RPMI + 10 % FBS and 50 ng ml^−1^ M-CSF and incubated overnight.

### Ethics statement

Blood donations for subsequent isolation of PBMCs were obtained from healthy donors after written, informed consent, in accordance with the Declaration of Helsinki. All protocols were approved by the Ethics Committee of the University Hospital Jena (permission number 2207-01/08).

### Co-incubation of *C. parapsilosis* with monocyte-derived macrophages (MDMs)

Overnight cultures of yeast cells were harvested by centrifugation at 5,000 rpm/4 °C for 5 min and cells were washed three times in ice-cold H_2_O.

### *P. aurantium* and macrophage killing assays

Yeast strains were grown overnight in YPD medium at 30°C and 180 rpm, harvested by centrifugation, and counted in a CASY® TT Cell Counter (OLS Bio, Bremen, Germany). Amoebae were grown to confluency in PB, harvested by scraping, and counted. Yeast cells were co-incubated with amoebae or macrophages in 96-well plates at MOIs of 10 and 1, respectively, and incubated at 22°C or 37°C/5% CO_2_, respectively, for 3 hours. Yeast cells surviving the amoeba predation were collected by vigorous pipetting and plated on YPD agar. For macrophage killing assays, yeast cells were added to macrophages in 96-well plates at an MOI of 1 (killing assays) or in 6-well plates at an MOI of 10 (isolation of total RNA) with RPMI medium containing L-glutamine. Media control wells for each time point were included, where the yeast cells were incubated in RPMI alone without macrophages. Plates were incubated at 37°C in an atmosphere with 5% CO_2_. Cells surviving the macrophage killing were first collected from the supernatant, then, intracellular survivors were obtained after lysis of macrophages with 0.5 % Triton™-X-100 for 15 min. The number of survived yeast cells was calculated as a percentage of CFUs compared to the inoculum. Data are based on three biological and six technical replicates (*P. aurantium*) and six different anonymous donors with 6 technical replicates (macrophages), respectively.

## Acknowledgements

This work was supported by grants of the European Social Fund ESF “Europe for Thuringia” (2015FGR0097 to FH and 2016FGR0053 to JL) and a grant from the German Research Foundation (DFG, HI 1574/2-1). SR was supported by a fellowship of the DFG funded excellence graduate school “Jena School of Microbial Communication“—JSMC. This work was also supported by the DFG CRC/Transregio 124 “Pathogenic fungi and their human host: Networks of interaction” subproject INF (TW). MS was supported within the DFG priority program SPP1580 ‘Intracellular compartments as places of host-pathogen interactions’ (Hu 528/17–1). AG was funded by GINOP-2.3.2-15-2016-00035, by GINOP-2.3.3-15-2016-00006 and by NKFIH K123952.

## Author contributions

SR performed most experimental work with input from JLS, RT, and MS. FH, AG, SB, and GP supervised the experimental work. Bioinformatic processing and analysis of RNA-Seq data was performed by JL and TW. SR and FH wrote the paper. All authors analyzed the data and commented on the manuscript.

## Declaration of Interests

The authors declare no competing interests.

## References

1. Dadar M, Tiwari R, Karthik K, Chakraborty S, Shahali Y, Dhama K. *Candida albicans* - Biology, molecular characterization, pathogenicity, and advances in diagnosis and control - An update. Microb Pathogen. 2018;117:128–38. Epub 2018/02/20.

2. Robinson HA, Pinharanda A, Bensasson D. Summer temperature can predict the distribution of wild yeast populations. Ecol Evol. 2016;6(4):1236–50.

3. Maganti H, Bartfai D, Xu J. Ecological structuring of yeasts associated with trees around Hamilton, Ontario, Canada. FEMS Yeast Res. 2012;12(1):9–19.

4. Bensasson D, Dicks J, Ludwig JM, Bond CJ, Elliston A, Roberts IN, et al. Diverse Lineages of *Candida albicans* Live on Old Oaks. Genetics. 2019;211(1):277.

5. Greppi A, Krych Ł, Costantini A, Rantsiou K, Hounhouigan DJ, Arneborg N, et al. Phytase-producing capacity of yeasts isolated from traditional African fermented food products and PHYPk gene expression of *Pichia kudriavzevii* strains. Int J Food Microbiol. 2015;205:81–9.

6. Morrison-Whittle P, Lee SA, Fedrizzi B, Goddard MR. Co-evolution as Tool for Diversifying Flavor and Aroma Profiles of Wines. Front Microbiol. 2018;9(910).

7. Erwig LP, Gow NA. Interactions of fungal pathogens with phagocytes. Nat Rev Microbiol. 2016;14(3):163–76.

8. Seider K, Heyken A, Luttich A, Miramon P, Hube B. Interaction of pathogenic yeasts with phagocytes: survival, persistence and escape. Curr Opin Microbiol. 2010;13(4):392–400. Epub 2010/07/16.

9. Uwamahoro N, Verma-Gaur J, Shen H-H, Qu Y, Lewis R, Lu J, et al. The Pathogen *Candida albicans* Hijacks Pyroptosis for Escape from Macrophages. MBio. 2014;5(2).

10. Wellington M, Koselny K, Sutterwala FS, Krysan DJ. *Candida albicans* triggers NLRP3-mediated pyroptosis in macrophages. Eukaryot Cell. 2014;13(2):329–40. Epub 2014/01/01.

11. Kasper L, Konig A, Koenig PA, Gresnigt MS, Westman J, Drummond RA, et al. The fungal peptide toxin Candidalysin activates the NLRP3 inflammasome and causes cytolysis in mononuclear phagocytes. Nat Commun. 2018;9(1):018–06607.

12. Toth R, Toth A, Papp C, Jankovics F, Vagvolgyi C, Alonso MF, et al. Kinetic studies of *Candida parapsilosis* phagocytosis by macrophages and detection of intracellular survival mechanisms. Front Microbiol. 2014;5:633. Epub 2014/12/06.

13. Linden JR, Maccani MA, Laforce-Nesbitt SS, Bliss JM. High Efficiency Opsonin-Independent Phagocytosis of *Candida parapsilosis* by Human Neutrophils. Med Mycol. 2010;48(2):10.1080/13693780903164566.

14. Sasada M, Johnston RB, Jr. Macrophage microbicidal activity. Correlation between phagocytosis-associated oxidative metabolism and the killing of *Candida* by macrophages. J Exp Med. 1980;152(1):85–98.

15. Kaur R, Ma B, Cormack BP. A family of glycosylphosphatidylinositol-linked aspartyl proteases is required for virulence of *Candida glabrata*. Proceed Natl Acad Sci USA. 2007;104(18):7628–33. Epub 2007/04/26.

16. Seider K, Brunke S, Schild L, Jablonowski N, Wilson D, Majer O, et al. The facultative intracellular pathogen *Candida glabrata* subverts macrophage cytokine production and phagolysosome maturation. J Immunol. 2011;187(6):3072–86.

17. Pesole G, Lotti M, Alberghina L, Saccone C. Evolutionary origin of nonuniversal CUG_Ser_ codon in some *Candida* species as inferred from a molecular phylogeny. Genetics. 1995;141(3):903–7.

18. Gabaldon T, Martin T, Marcet-Houben M, Durrens P, Bolotin-Fukuhara M, Lespinet O, et al. Comparative genomics of emerging pathogens in the *Candida glabrata* clade. BMC Genomics. 2013;14(623):1471–2164.

19. Turner SA, Butler G. The *Candida* pathogenic species complex. Cold Spring Harb Perspect Med. 2014;4(9).

20. Gabaldon T, Fairhead C. Genomes shed light on the secret life of *Candida glabrata*: not so asexual, not so commensal. Curr Genet. 2019;65(1):93–8.

21. Casadevall A, Fu MS, Guimaraes AJ, Albuquerque P. The ‘Amoeboid Predator-Fungal Animal Virulence’ Hypothesis. J Fungi. 2019;5(1).

22. Steenbergen JN, Nosanchuk JD, Malliaris SD, Casadevall A. *Cryptococcus neoformans* virulence is enhanced after growth in the genetically malleable host *Dictyostelium discoideum*. Infect Immun. 2003;71(9):4862–72.

23. Steenbergen JN, Shuman HA, Casadevall A. *Cryptococcus neoformans* interactions with amoebae suggest an explanation for its virulence and intracellular pathogenic strategy in macrophages. Proceed Natl Acad Sci USA. 2001;98(26):15245–50.

24. Van Waeyenberghe L, Bare J, Pasmans F, Claeys M, Bert W, Haesebrouck F, et al. Interaction of *Aspergillus fumigatus* conidia with *Acanthamoeba castellanii* parallels macrophage-fungus interactions. Environ Microbiol Rep. 2013;5(6):819–24.

25. Hillmann F, Novohradska S, Mattern DJ, Forberger T, Heinekamp T, Westermann M, et al. Virulence determinants of the human pathogenic fungus *Aspergillus fumigatus* protect against soil amoeba predation. Environ Microbiol. 2015;17(8):2858–69.

26. Koller B, Schramm C, Siebert S, Triebel J, Deland E, Pfefferkorn AM, et al. *Dictyostelium discoideum* as a Novel Host System to Study the Interaction between Phagocytes and Yeasts. Front Microbiol. 2016;7(1665).

27. Aguilar M, Lado C, Spiegel FW. Protostelids from deciduous forests: first data from southwestern Europe. Mycol Res. 2007;111(Pt 7):863–72. Epub 2007/08/08.

28. Ndiritu GG, Stephenson SL, Spiegel FW. First records and microhabitat assessment of protostelids in the Aberdare Region, Central Kenya. J Eukaryot Microbiol. 2009;56(2):148–58. Epub 2009/05/22.

29. Shadwick JD, Stephenson SL, Spiegel FW. Distribution and ecology of protostelids in Great Smoky Mountains National Park. Mycologia. 2009;101(3):320–8. Epub 2009/06/20.

30. Zahn G, Stephenson SL, Spiegel FW. Ecological distribution of protosteloid amoebae in New Zealand. PeerJ. 2014;2:e296. Epub 2014/04/02.

31. Hillmann F, Forbes G, Novohradska S, Ferling I, Riege K, Groth M, et al. Multiple Roots of Fruiting Body Formation in Amoebozoa. Genome Biol Evol. 2018;10(2):591–606. Epub 2018/01/30.

32. Radosa S, Ferling I, Sprague JL, Westermann M, Hillmann F. The different morphologies of yeast and filamentous fungi trigger distinct killing and feeding mechanisms in a fungivorous amoeba. Environ Microbiol. 2019;13(10):1462–2920.

33. Bochman ML, Schwacha A. The Mcm Complex: Unwinding the Mechanism of a Replicative Helicase. Microbiol Mol Biol Rev. 2009;73(4):652–83.

34. Navarathna DH, Lionakis MS, Lizak MJ, Munasinghe J, Nickerson KW, Roberts DD. Urea amidolyase (DUR1,2) contributes to virulence and kidney pathogenesis of *Candida albicans*. PloS one. 2012;7(10):e48475. Epub 2012/11/13.

35. Skrzypek MS, Binkley J, Binkley G, Miyasato SR, Simison M, Sherlock G. The Candida Genome Database (CGD): incorporation of Assembly 22, systematic identifiers and visualization of high throughput sequencing data. Nucleic Acids Res. 2017;45(D1):D592–D6. Epub 2016/10/13.

36. Weissman Z, Berdicevsky I, Cavari BZ, Kornitzer D. The high copper tolerance of *Candida albicans* is mediated by a P-type ATPase. Proceed Natl Acad Sci USA. 2000;97(7):3520–5. Epub 2000/03/29.

37. Srinivasa K, Kim NR, Kim J, Kim M, Bae JY, Jeong W, et al. Characterization of a putative thioredoxin peroxidase prx1 of *Candida albicans*. Mol Cells. 2012;33(3):301–7. Epub 2012/03/07.

38. Brussow H. Bacteria between protists and phages: from antipredation strategies to the evolution of pathogenicity. Mol Microbiol. 2007;65(3):583–9.

39. Old KM, Darbyshire JF. Soil fungi as food for giant amoebae. Soil Biol and Biochem. 1978;10(2):93–100.

40. Old KM. Giant soil amoebae cause perforation of conidia of *Cochliobolus sativus*. Transact Brit Mycol Soc. 1977;68(2):277–81.

41. Chakraborty S, Old KM, Warcup JH. Amoebae from a take-all suppressive soil which feed on *Gaeumannomyces graminis tritici* and other soil fungi. Soil Biol Biochem. 1983;15(1):17–24.

42. Spiegel FW, Shadwick LL, Ndiritu GG, Brown MW, Aguilar M, Shadwick JD. Protosteloid Amoebae (Protosteliida, Protosporangiida, Cavosteliida, Schizoplasmodiida, Fractoviteliida, and Sporocarpic Members of Vannellida, Centramoebida, and Pellitida). In: Archibald JM, Simpson AGB, Slamovits CH, Margulis L, Melkonian M, Chapman DJ, et al., editors. Handbook of the Protists. Cham: Springer International Publishing; 2017. p. 1–38.

43. Kang S, Tice AK, Spiegel FW, Silberman JD, Panek T, Cepicka I, et al. Between a Pod and a Hard Test: The Deep Evolution of Amoebae. Mol Biol Evol. 2017;34(9):2258–70.

44. Shadwick LL, Spiegel FW, Shadwick JDL, Brown MW, Silberman JD. Eumycetozoa = Amoebozoa?: SSUrDNA Phylogeny of Protosteloid Slime Molds and Its Significance for the Amoebozoan Supergroup. PloS one. 2009;4(8):e6754.

45. Shadwick JDL, Silberman JD, Spiegel FW. Variation in the SSUrDNA of the Genus Protostelium Leads to a New Phylogenetic Understanding of the Genus and of the Species Concept for *Protostelium mycophaga* (Protosteliida, Amoebozoa). J Eukaryot Microbiol. 2018;65(3):331–44.

46. Kasper L, Seider K, Hube B. Intracellular survival of *Candida glabrata* in macrophages: immune evasion and persistence. FEMS Yeast Research. 2015;15(5):fov042. Epub 2015/06/13.

47. Seider K, Gerwien F, Kasper L, Allert S, Brunke S, Jablonowski N, et al. Immune evasion, stress resistance, and efficient nutrient acquisition are crucial for intracellular survival of *Candida glabrata* within macrophages. Eukaryot Cell. 2014;13(1):170–83.

48. Lengeler KB, Tielker D, Ernst JF. Protein-O-mannosyltransferases in virulence and development. Cell Mol Life Sci. 2007;65(4):528.

49. Peltroche-Llacsahuanga H, Goyard S, d’Enfert C, Prill SK, Ernst JF. Protein O-mannosyltransferase isoforms regulate biofilm formation in *Candida albicans*. Antimicrob Agents Chemother. 2006;50(10):3488–91.

50. Hall RA, Bates S, Lenardon MD, Maccallum DM, Wagener J, Lowman DW, et al. The Mnn2 mannosyltransferase family modulates mannoprotein fibril length, immune recognition and virulence of *Candida albicans*. PLoS Pathog. 2013;9(4):25.

51. Hall RA, Gow NAR. Mannosylation in *Candida albicans*: role in cell wall function and immune recognition. Mol Microbiol. 2013;90(6):1147–61.

52. Bain JM, Louw J, Lewis LE, Okai B, Walls CA, Ballou ER, et al. *Candida albicans* hypha formation and mannan masking of beta-glucan inhibit macrophage phagosome maturation. MBio. 2014;5(6):01874–14.

53. McKenzie CG, Koser U, Lewis LE, Bain JM, Mora-Montes HM, Barker RN, et al. Contribution of *Candida albicans* cell wall components to recognition by and escape from murine macrophages. Infect Immun. 2010;78(4):1650–8.

54. Cote P, Hogues H, Whiteway M. Transcriptional analysis of the *Candida albicans* cell cycle. Mol Biol Cell. 2009;20(14):3363–73.

55. Wolschendorf F, Ackart D, Shrestha TB, Hascall-Dove L, Nolan S, Lamichhane G, et al. Copper resistance is essential for virulence of *Mycobacterium tuberculosis*. Proceed Natl Acad Sci USA. 2011;108(4):1621–6.

56. Hao X, Luthje F, Ronn R, German NA, Li X, Huang F, et al. A role for copper in protozoan grazing - two billion years selecting for bacterial copper resistance. Mol Microbiol. 2016;102(4):628–41.

57. Ding C, Festa RA, Chen Y-L, Espart A, Palacios Ò, Espín J, et al. *Cryptococcus neoformans* copper detoxification machinery is critical for fungal virulence. Cell Host Microbe. 2013;13(3):265–76.

58. Wiemann P, Perevitsky A, Lim FY, Shadkchan Y, Knox BP, Landero Figueora JA, et al. *Aspergillus fumigatus* Copper Export Machinery and Reactive Oxygen Intermediate Defense Counter Host Copper-Mediated Oxidative Antimicrobial Offense. Cell Rep. 2017;19(5):1008–21.

59. Novohradska S, Ferling I, Hillmann F. Exploring Virulence Determinants of Filamentous Fungal Pathogens through Interactions with Soil Amoebae. Front Cell Infect Microbiol. 2017;7:497. Epub 2017/12/21.

60. Douglas LM, Konopka JB. Plasma membrane architecture protects *Candida albicans* from killing by copper. PLoS Genet. 2019;15(1):e1007911–e.

61. Li CX, Gleason JE, Zhang SX, Bruno VM, Cormack BP, Culotta VC. *Candida albicans* adapts to host copper during infection by swapping metal cofactors for superoxide dismutase. Proceed Natl Acad Sci USA. 2015;112(38):E5336–E42.

62. Lorenz MC, Bender JA, Fink GR. Transcriptional Response of *Candida albicans* upon Internalization by Macrophages. Eukaryotic Cell. 2004;3(5):1076–87.

63. Rubin-Bejerano I, Fraser I, Grisafi P, Fink GR. Phagocytosis by neutrophils induces an amino acid deprivation response in *Saccharomyces cerevisiae* and *Candida albicans*. Proceed Natl Acad Sci USA. 2003;100(19):11007–12.

64. Miramón P, Dunker C, Windecker H, Bohovych IM, Brown AJP, Kurzai O, et al. Cellular Responses of *Candida albicans* to Phagocytosis and the Extracellular Activities of Neutrophils Are Critical to Counteract Carbohydrate Starvation, Oxidative and Nitrosative Stress. PloS one. 2012;7(12):e52850.

65. Chang EC, Kosman DJ. O_2_-dependent methionine auxotrophy in Cu,Zn superoxide dismutase-deficient mutants of *Saccharomyces cerevisiae*. J Bacteriol. 1990;172(4):1840–5.

66. Culotta VC, Klomp LW, Strain J, Casareno RL, Krems B, Gitlin JD. The copper chaperone for superoxide dismutase. J Biol Chem. 1997;272(38):23469–72.

67. Andrew S. FastQC: a quality control tool for high throughput sequence data. Available online at: http://www.bioinformatics.babraham.ac.uk/projects/fastqc2010.

68. Bolger AM, Lohse M, Usadel B. Trimmomatic: a flexible trimmer for Illumina sequence data. Bioinformatics. 2014;30(15):2114–20.

69. Kim D, Pertea G, Trapnell C, Pimentel H, Kelley R, Salzberg SL. TopHat2: accurate alignment of transcriptomes in the presence of insertions, deletions and gene fusions. Genome Biol. 2013;14(4):R36.

70. Robinson MD, McCarthy DJ, Smyth GK. edgeR: a Bioconductor package for differential expression analysis of digital gene expression data. Bioinformatics. 2010;26(1):139–40.

71. Mi H, Huang X, Muruganujan A, Tang H, Mills C, Kang D, et al. PANTHER version 11: expanded annotation data from Gene Ontology and Reactome pathways, and data analysis tool enhancements. Nucleic Acids Res. 2017;45(D1):D183–D9.

72. Noble SM, Johnson AD. Strains and strategies for large-scale gene deletion studies of the diploid human fungal pathogen *Candida albicans*. Eukaryot Cell. 2005;4(2):298–309.

73. Holland LM, Schroder MS, Turner SA, Taff H, Andes D, Grozer Z, et al. Comparative phenotypic analysis of the major fungal pathogens *Candida parapsilosis* and *Candida albicans*. PLoS Pathog. 2014;10(9):e1004365. Epub 2014/09/19.

74. Gerami-Nejad M, Zacchi LF, McClellan M, Matter K, Berman J. Shuttle vectors for facile gap repair cloning and integration into a neutral locus in *Candida albicans*. Microbiol. 2013;159(Pt 3):565–79. Epub 2013/01/12.

