## Supplementary material for "Phagocytic predation by the fungivorous amoeba *Protostelium aurantium* targets metal ion and redox homeostasis": Radosa_etal_2019_SFigures.pdf

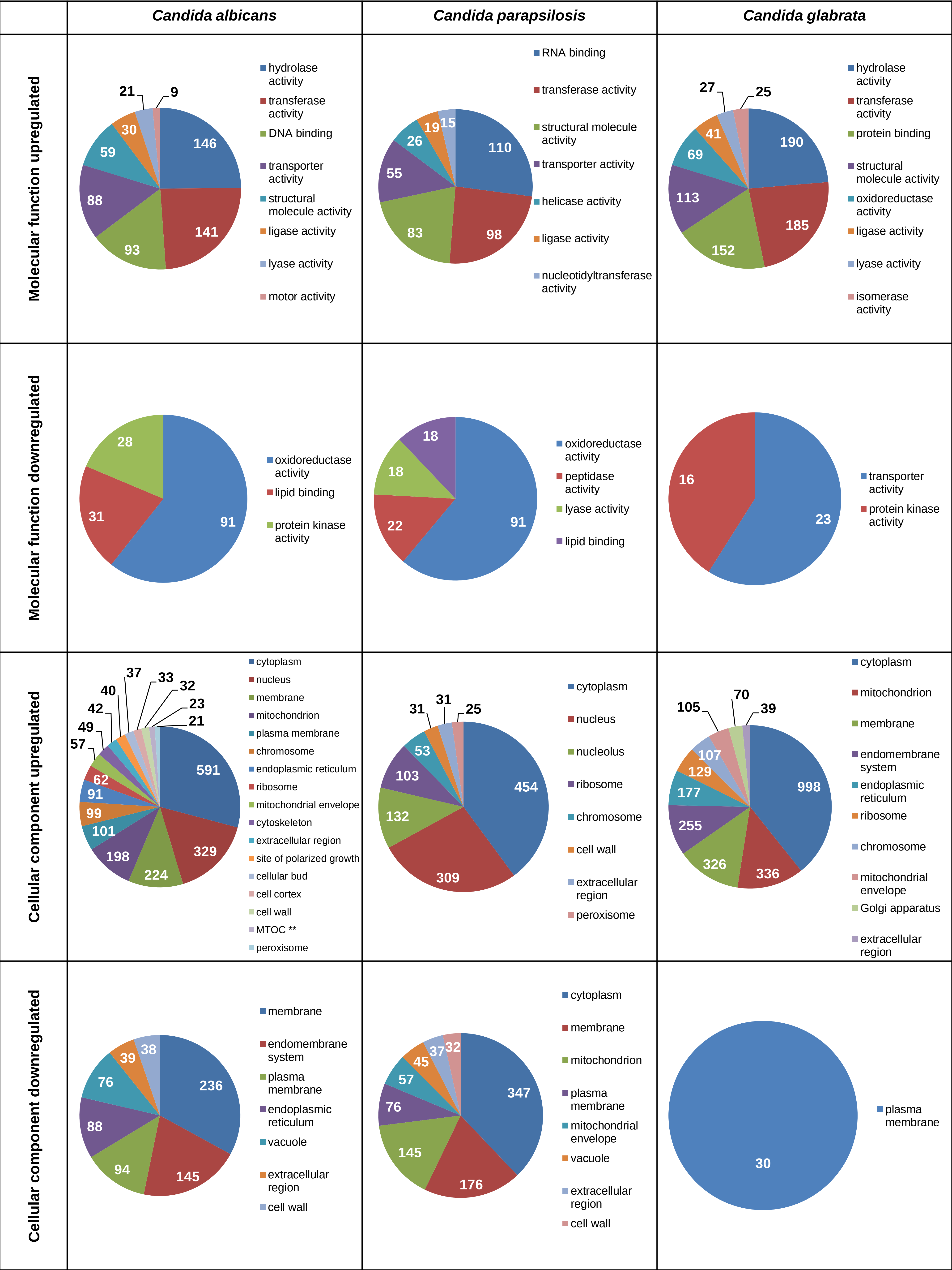

Candida albicans
Candida parapsilosis
Candida glabrata
Molecular function upregulated
146
141
93
88
59
30
21
9
hydrolase activity
transferase activity
DNA binding
transporter activity
structural molecule activity
ligase activity
lyase activity
motor activity
110
98
83
55
26
19
15
RNA binding
transferase activity
structural molecule activity
transporter activity
helicase activity
ligase activity
nucleotidyltransferase activity
190
185
152
113
69
41
27
25
hydrolase activity
transferase activity
protein binding
structural molecule activity
oxidoreductase activity
ligase activity
lyase activity
isomerase activity
Molecular function downregulated
91
31
28
oxidoreductase activity
lipid binding
protein kinase activity
91
22
18
18
oxidoreductase activity
peptidase activity
lyase activity
lipid binding
23
16
transporter activity
protein kinase activity
Cellular component upregulated
591
329
224
198
101
99
91
62
57
49
42
40
37
33
32
23
21
cytoplasm
nucleus
membrane
mitochondrion
plasma membrane
chromosome
endoplasmic reticulum
ribosome
mitochondrial envelope
cytoskeleton
extracellular region
site of polarized growth
cellular bud
cell cortex
cell wall
MTOC **
peroxisome
454
309
132
103
53
31
31
25
cytoplasm
nucleus
nucleolus
ribosome
chromosome
cell wall
extracellular region
peroxisome
998
336
326
255
177
129
107
105
70
39
cytoplasm
mitochondrion
membrane
endomembrane system
endoplasmic reticulum
ribosome
chromosome
mitochondrial envelope
Golgi apparatus
extracellular region
Cellular component downregulated
236
145
94
88
76
39
38
membrane
endomembrane system
plasma membrane
endoplasmic reticulum
vacuole
extracellular region
cell wall
347
176
145
76
57
45
37
32
cytoplasm
membrane
mitochondrion
plasma membrane
mitochondrial envelope
vacuole
extracellular region
cell wall
30
plasma membrane

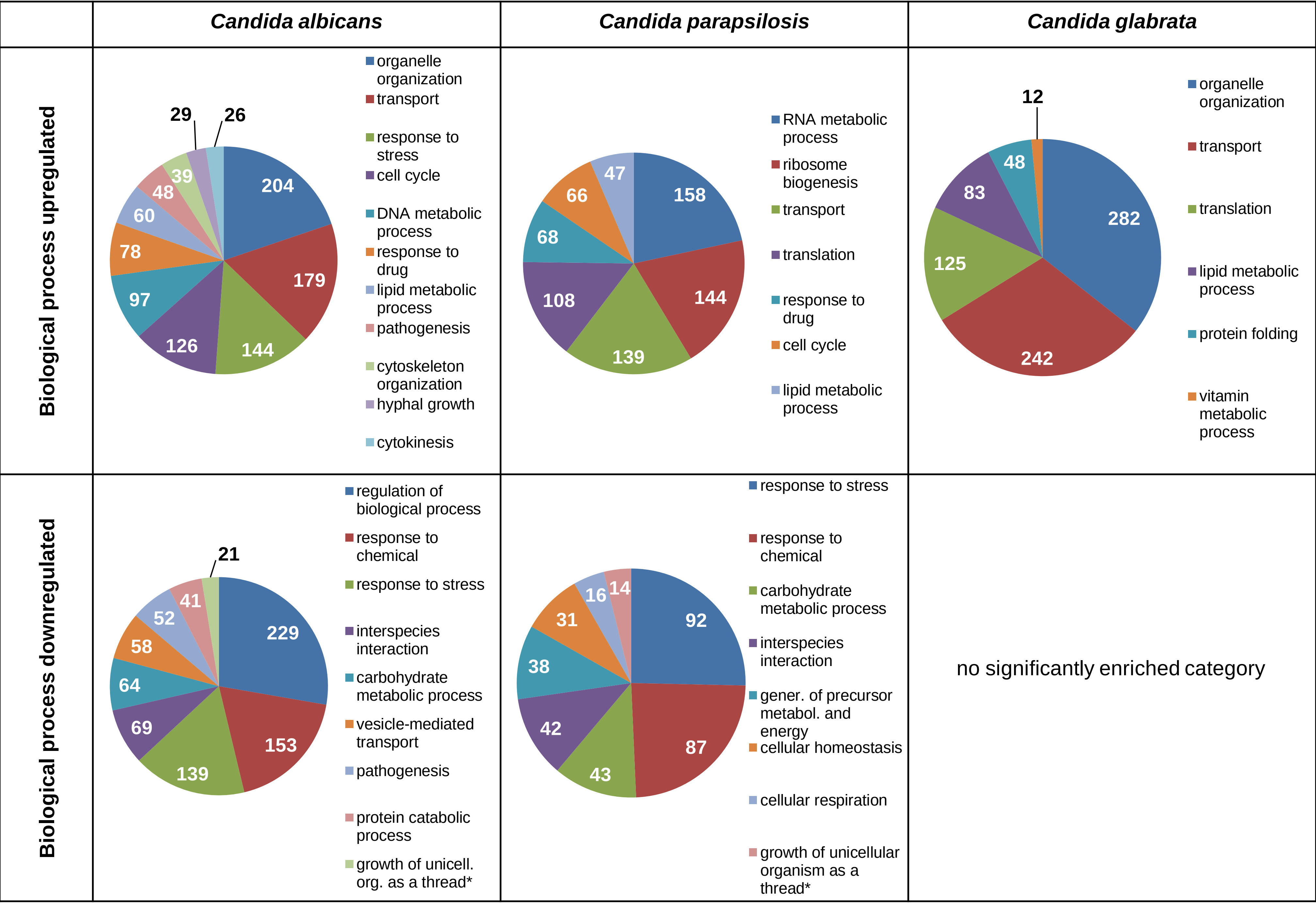


**Fig. S1: Cluster analysis of differentially expressed genes in *C. albicans, C. parapsilosis* and *C. glabrata* upon predation by *P. aurantium*.** Positively and negatively regulated genes we assign to categories according the molecular function (MF), cellular component (CC) and biological process (BP) using an on-line tool Slim Mapper at Candida Genome Database. Enriched GO terms, displayed in the pie charts, we determined by the Benjamini-Hochberg corrected *p*-value. The numbers indicate the number of DEG assign to the particular cluster. A full list of genes and enriched GO terms in provided in Suppl. Dataset IV.

* growth of unicellular organism as a thread to attached cells (BP:GO:0070783)

** microtubule organizing center (CC:GO:0005815)

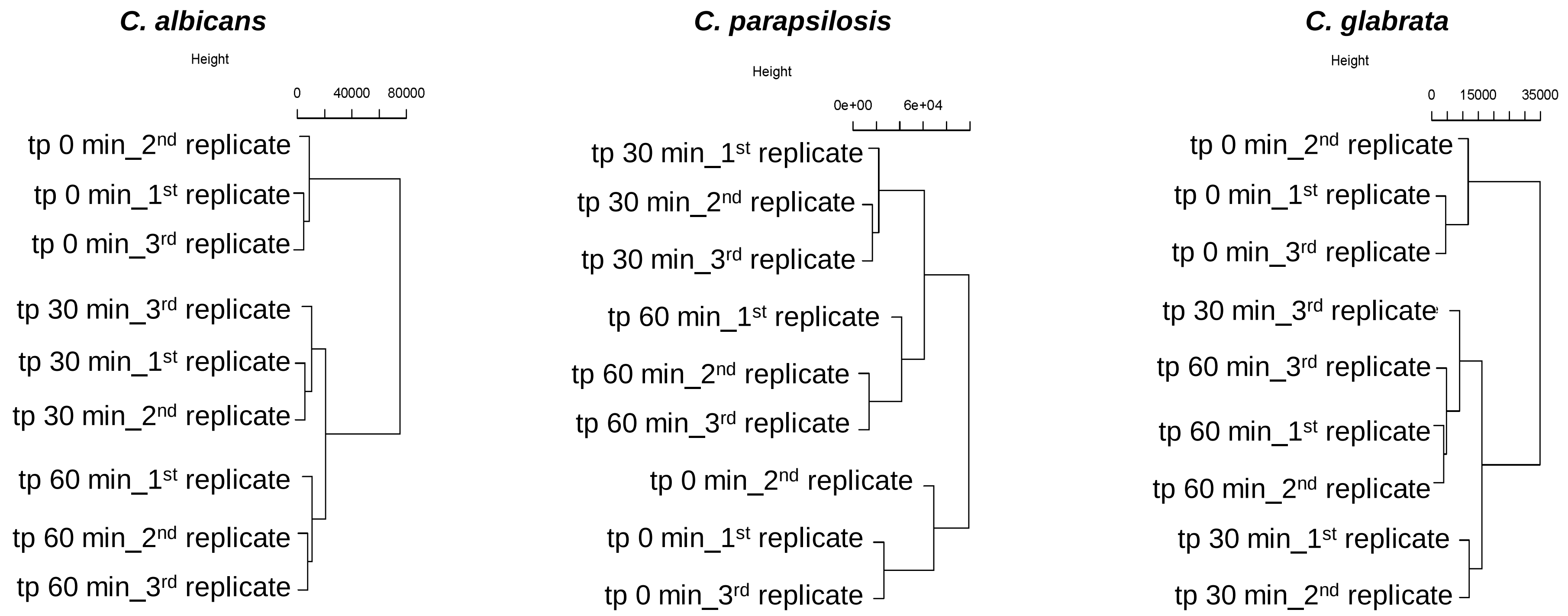


**Fig. S2: Cluster dendrogram of three *Candida* species transcription profiles in a response to amoeba predation.**

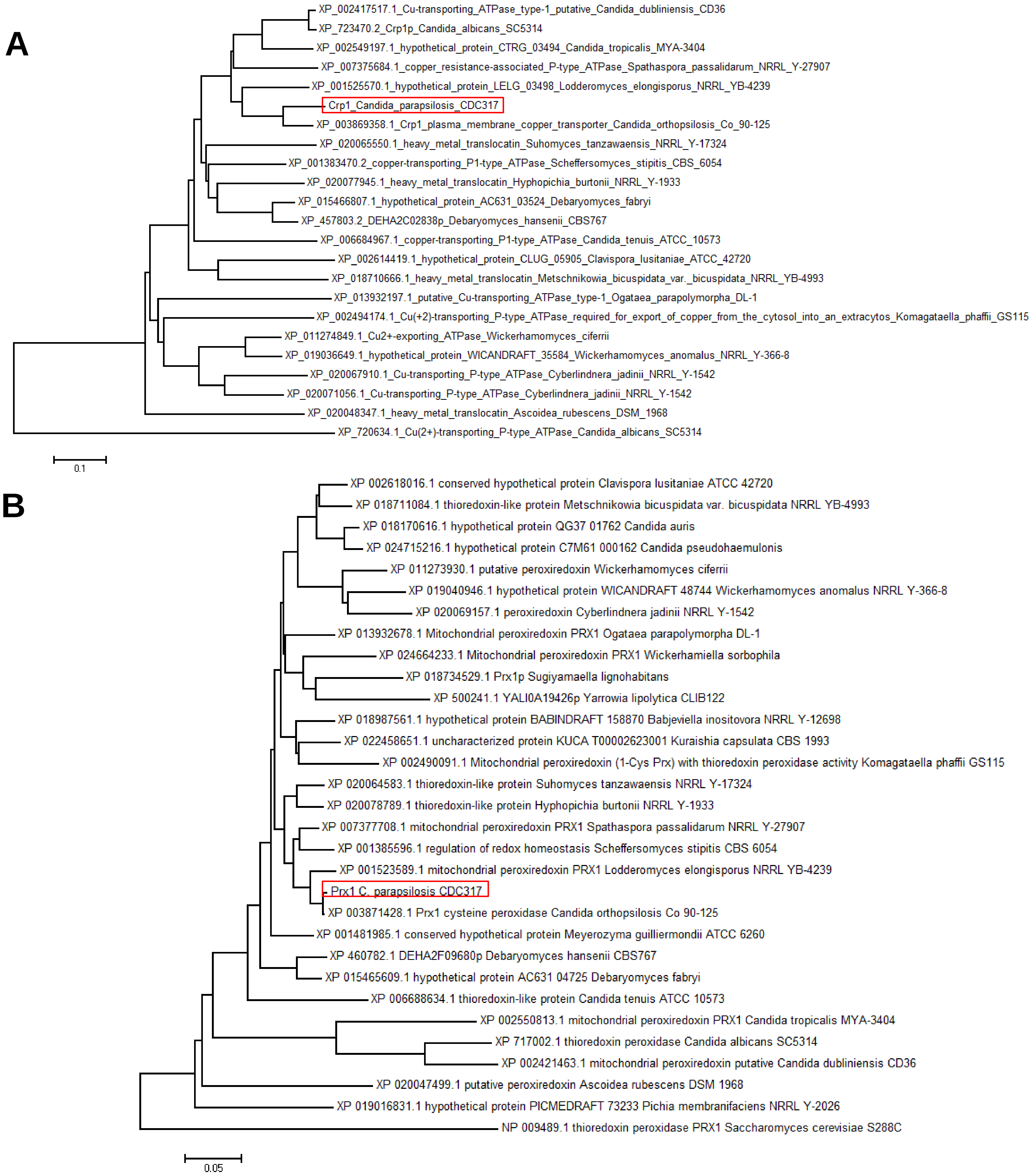


**Fig. S3: Neighbor joing trees for Crp1 (A) and Prx1 (B) homologous proteins within the *Saccharomycotina*.** Trees were based on phylogeny reconstruction and generated from Muscle Alignments (Edgar, 2004) using MEGA 6 software (Tamura et al., 2013) at the following settings: Gap Open, -2.9; Gap Extend, 0; Hydrophobicity Multiplier, 1.2 for the alignment. The Ccc2 protein of *C. albicans* and the Prx1 protein from *S. cerevisiae* were used as outgroups. Red boxes mark the Crp1 and Prx1 proteins of *C. parapsilosis.*

Edgar RC. 2004. MUSCLE: multiple sequence alignment with high accuracy and high throughput. *Nucl Acids Res* 32:1792-1797.
Tamura K, Stecher G, Peterson D, Filipski A, and Kumar S. 2013. MEGA6: Molecular Evolutionary Genetics Analysis Version 6.0. *Mol Biol Evol* 30:2725-2729.

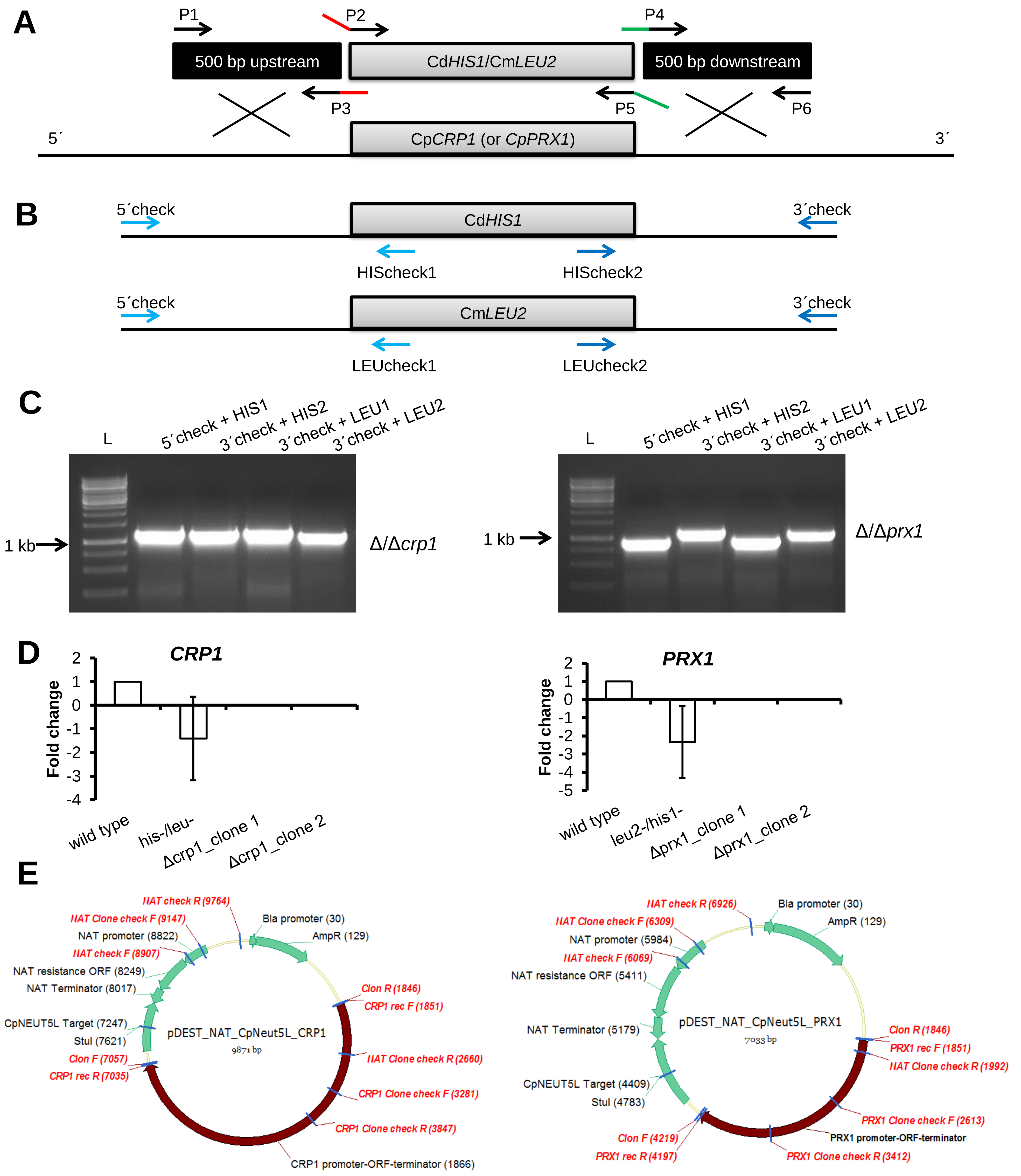


**Fig. S4: Construction of deletion mutants and complemented strains of *CRP1* and *PRX1* genes in *C. parapsilosis*.** (A) Scheme depicting generation of gene knockout strains. Both alleles of each gene were deleted by replacing them with selectable markers *C. dubliniensis HIS1* and *C. maltosa LEU2* via homologous recombination. Primer overlaps are shown in color. For each strain, two independent deletion mutants were created. Picture is modified from Holland *et al*., 2014. **B+C,** Correct integration of disruption cassette within the target locus was verified in 4 PCR reactions detecting both recombinant upstream and downstream loci for each of the alleles. **D**, The absence of functional genes was documented by the absence of detectable transcript in Real-Time PCR. Both mutants displayed no expression of the deletion targets when testing two independent clones. **E**, Plasmid maps for the generation of complemented strains. Primer binding sites are indicated in red.
